# *Yersinia pestis* lipoprotein SlyB promotes plague pathogenesis via envelope stress tolerance

**DOI:** 10.64898/2026.06.08.730324

**Authors:** Pierre Lê-Bury, Emelyne Bougit, Sébastien Bontemps-Gallo, Guillem Mas Fiol, Cyril Savin, Van-Son Nguyen, Julien Madej, Rémi Beau, Caroline Buscail, Nicolas Bouladoux, Friederike Jönsson, COSIPOP Study group, Marion Lemarignier, Mara Emilia Carloni, Anne Derbise, Christian Demeure, Florent Sebbane, Han Remaut, Javier Pizarro-Cerda, Olivier Dussurget

## Abstract

*Yersinia pestis*, the etiological agent of plague, persists in an enzootic cycle involving mammals and fleas, requiring constant outer membrane (OM) adaptation to disparate host environments. One such pathway involves the glycine zipper 2TM domain-containing protein SlyB, a central component of the OM stress response and PhoPQ virulence pathway. While the OM is critical for virulence, the role of the OM lipoprotein SlyB in *Y. pestis* ecology and pathogenesis remains unknown. We show by phylogenetic analyses that *slyB* paralogs expanded in environmental bacteria, whereas the canonical *slyB* gene was under negative selective pressure during *Y. pestis* speciation from *Yersinia pseudotuberculosis*. Using rodent and flea infection models recapitulating *Y. pestis* natural history, we demonstrate that SlyB is specifically required to resist the mammalian immune system at 37°C, including neutrophil-mediated antimicrobial activity during lymph node colonization, but is dispensable in septicemic plague in rodents. Strikingly, SlyB is not required for flea colonization and resistance to the antimicrobial-peptide-based immunity of arthropods at lower temperatures. SlyB-dependent OM stress tolerance reveals a mechanism by which *Y. pestis* establishes bubonic plague, in line with its critical lipopolysaccharide structural switch. Our findings identify SlyB as an evolutionarily fine-tuned component of the *Y. pestis* envelope that mediates immune escape upon infection of mammalian hosts through maintenance of structural integrity.

## Introduction

*Yersinia pestis* is the etiological agent of plague, a highly lethal disease which claimed hundreds of millions of lives during three major pandemics throughout history^1^. It is a recent clone originating from the enteropathogenic bacterium *Yersinia pseudotuberculosis*, which diverged less than 6,000 years ago and drastically changed lifestyle through a series of gene acquisitions and gene losses^2^. It is known to persists through a complex enzootic cycle involving numerous rodents as primary reservoirs and fleas as vectors^3^. Mammalian infection usually occurs through injection of *Y. pestis* in the dermis upon an infected flea bite^4^. Bacteria quickly migrate through lymph flow from the site of injection to the draining lymph node (dLN), where they replicate and prevent immune system activity via diverse virulence factors, mainly targeting neutrophils and monocytes^5–7^. After disruption of lymph node and blood vessel’s structure^8^, and hematogenous dissemination, *Y. pestis* replicates in the liver, spleen and blood leading to a high-level bacteremia (10^6^ to 10^7^ bacteria per mL) and quickly to death. Vector infection occurs during a blood meal in this short bacteremic phase^9^. Transmission from infected vector to mammals proceeds via bacterial regurgitation through distinct dynamics: an "early-phase" transmission, a "late-phase" transmission resulting from proventricular blockage by a biofilm; and an intermediate state of partial blockage^10^. Throughout this cycle, *Y. pestis* needs to adapt to different ecological niches, host constraints and temperatures: biofilms are formed around 21°C in the flea proventriculus; bacteria grown at low temperature are inoculated into the mammalian host dermis, then bacteria replicate in different organs at 37°C; and finally back to the flea at lower-temperature after a blood meal.

The bacterial envelope is a crucial interface during host-pathogen interactions, and numerous studies explored the impact of structural envelope components on *Y. pestis* virulence. The abundant outer membrane protein (OMP) OmpA and the Braun lipoprotein Lpp are required for outer membrane (OM) attachment to peptidoglycan and plague pathogenesis^11–13^; LPS modifications serve both vector and host infection, as 4-amino-4-deoxy-L-arabinose (Ara4N) addition to lipid A could allow to counter antimicrobial peptides-based arthropod immune defense at flea temperature^14,15^, while switch from hexa- to tetra-acylated LPS leads to immune escape when transitioning to mammalian host temperature^16–18^. Envelope also plays an important role for environmental sensing and pathogenesis, as two-components systems (TCS) such as PhoPQ and EnvZ/OmpR regulate some of the above-mentioned mechanisms *in vitro*, and are required for successful host infection^19–21^. Finally, the most important *Y. pestis* virulence determinants interact with the bacterial envelope, such as the type three secretion system (T3SS) spanning through inner membrane, periplasmic space and outer membrane^22^; the plasminogen activator Pla, a protease inserted in the OM whose activity depends on LPS conformation^23^; *Y. pestis* most abundant OMP, the attachment-invasion locus Ail, involved in serum resistance and OM structural integrity^24–26^; the fimbriae PsaA and the pseudocapsule Caf, attached to the envelope and involved in phagocytosis resistance or flea transmission^27–30^. These surface proteins also participate in cell adhesion facilitating *Yersinia* outer proteins (Yop) delivery by the T3SS^31^.

Recently the SlyB outer-membrane lipoprotein has been described as a new key player in bacterial envelope homeostasis, through its role in OM stress tolerance via OMP encapsulation in lipid nanodomains^32^. SlyB was described more than 30 years ago for the first time as a peptidoglycan-associated lipoprotein (PAL) cross-reacting protein and was first called PCP^33^. The alternative SlyB denomination originates from the proximity of *slyB* gene locus with the *slyA* gene^34^, which was initially erroneously identified as encoding a hemolysin in *Salmonella enterica*, hence called the salmolysin SlyA^35^. In fact, *slyA* (also called regulator of virulence A *rovA* in *Yersinia* species^36^) encodes a transcriptional regulator which induces expression of a hemolysin in *Escherichia coli* but not in *S. enterica*^34^. SlyB is well-conserved among diderm bacteria and is part of the core regulon of the PhoPQ TCS in several species^37^, which has been confirmed experimentally in *E. coli*^38^, *S. enterica*^39^, *Shigella flexneri*^40^, *Pseudomonas aeruginosa*^41^ and *Y. pestis*^37^. It was suggested that the broad conservation of SlyB was due to its activity as negative regulator of PhoPQ, as shown in *S. enterica*^37^. However, it is not the case in *E. coli* and the conserved negative regulation of PhoQ would rather come from the PhoPQ-regulated small peptide MgrB in *E. coli*, *S. enterica* and *Y. pestis*^42^. SlyB has been previously identified in transposon screens as a factor involved in intracellular survival of *E. coli* in macrophages^43,44^, a phenotype recently confirmed with *S. enterica*^32^. While several studies showed a role of SlyB in membrane stress tolerance^32,41,45^, implication of SlyB in host-pathogen interactions, especially for highly virulent pathogens such as *Y. pestis*, remains to be determined.

Taking advantage of the recently published structure of *E. coli* SlyB, here we decipher *Y. pestis* SlyB contribution to the natural ecological history of this pathogen. We show that SlyB was under evolutionary pressure during *Y. pestis* emergence, and that its function in stress tolerance is required in its natural mammalian host to counter neutrophil-mediated antimicrobial activity during the lymph node colonization step of bubonic plague, but neither for septicemic plague nor flea colonization at lower temperature. We propose that SlyB central role in membrane stress tolerance could have helped shape the delicate envelope equilibrium required for *Y. pestis* immune escape, high virulence and structural integrity.

## Results

### Evolutionary expansion and inactivation of *slyB* loci during *Yersinia* speciation

We identified 3 *slyB* paralogs in *Y. pestis* CO92 reference genome (NC_003143.1): the canonical *slyB* gene at locus YPO2373 (RefSeq locus YPO_RS12855) located near *slyA* (also called *rovA*); a paralog at locus YPO3646 (RefSeq locus YPO_RS19215); and a frameshifted paralog at RefSeq locus YPO_RS04730. Sequence alignment of canonical *E. coli* SlyB (SlyB*_Ec_*), *Y. pestis* canonical SlyB (SlyB*_Yp_*) and paralog 2, and the unframeshifted paralog 3 from *Y. pseudotuberculosis* showed conserved residues and functional structure such as the 2 transmembrane α-helix containing the glycine zipper (2TM Gly zipper) as well as the 3 phospholipid (PL)-binding residues described earlier^32^ (Fig. 1A). Amino acid (aa) sequences showed high homology, with 74.84%, 66.67%, and 61.94% identity between full-length SlyB_Ec_ to SlyB_Yp_, paralog 2, and paralog 3 from *Y. pseudotuberculosis*, respectively. Excluding the less conserved signal peptide increased sequence identity to 79.71%, 71.22%, and 66.67%, respectively. Comparison of the solved structure from SlyB_Ec_ and AlphaFold predictions showed a highly similar structure for SlyB*_Ec_* and SlyB*_Yp_* (Fig. 1B). Predictions of the noncanonical *Yersinia* paralogs show some structural variation at the C-terminus of the periplasmic domain, and a low prediction confidence in the 2TM Gly zipper transmembrane domains (Fig. 1B). To decipher the origin of these paralogs, SlyB orthologs were searched in the whole *Yersinia* genus (Supplementary Fig. 1, 2, Supplementary Table 1). The most ancestral *Yersinia entomophaga* and *Yersinia ruckeri* (insect and fish pathogens, respectively) and *Yersinia nurmii* encoded a unique canonical SlyB ortholog situated between the *anmK* and the *slyA* (*rovA*) gene, (Supplementary Fig. 3). The two other *slyB* paralogs were present in most of the other *Yersinia* species. The paralog 2 was usually situated between *dusB*, *fis* and a diguanylate cyclase on one end, and loci encoding cold-shock proteins CspA and CspG, transporters and the lipopolysaccharide-modifying protein locus *lpxP* on the other end, with other genes inserted in-between in some isolates. Paralog 3 was generally located between the hydrogenase *hyb* operon locus on one end and *lafU* (*motB*) and *motA* on the other end (Supplementary Fig. 3). In *Yersinia similis* and in the *Yersinia pseudotuberculosis* complex, this last paralog was situated next to *yjeO* and a DUF805 containing locus instead of the *hyb* operon. *Yersinia aldovae*, situated at the crossroad between these two clades, did not harbor the third paralog, suggesting it was deleted during genetic rearrangement of this region. We looked in genomes from closely related species in the *Yersiniceae* family such as *Serratia marcescens*, *Serratia aquatilis*, *Rahnella sikkimica* or *Chania multitudinisentens*, and these *slyB* paralogs were not found between *lpxP* and *fis*/*dusB* loci, neither near the *hyb* or *mot* operons. This suggests that paralogs 2 and 3 were acquired within the *Yersinia* genus after speciation from *Y. entomophaga* and *Y. ruckeri*. The third paralog was pseudogenized early in *Y. pestis* evolution, appearing frameshifted or absent in most of the recent *Y. pestis* isolates, with potential reversion mutations occurring in few clades (Supplementary Fig. 4, Supplementary Table 2).

**Figure 1.**
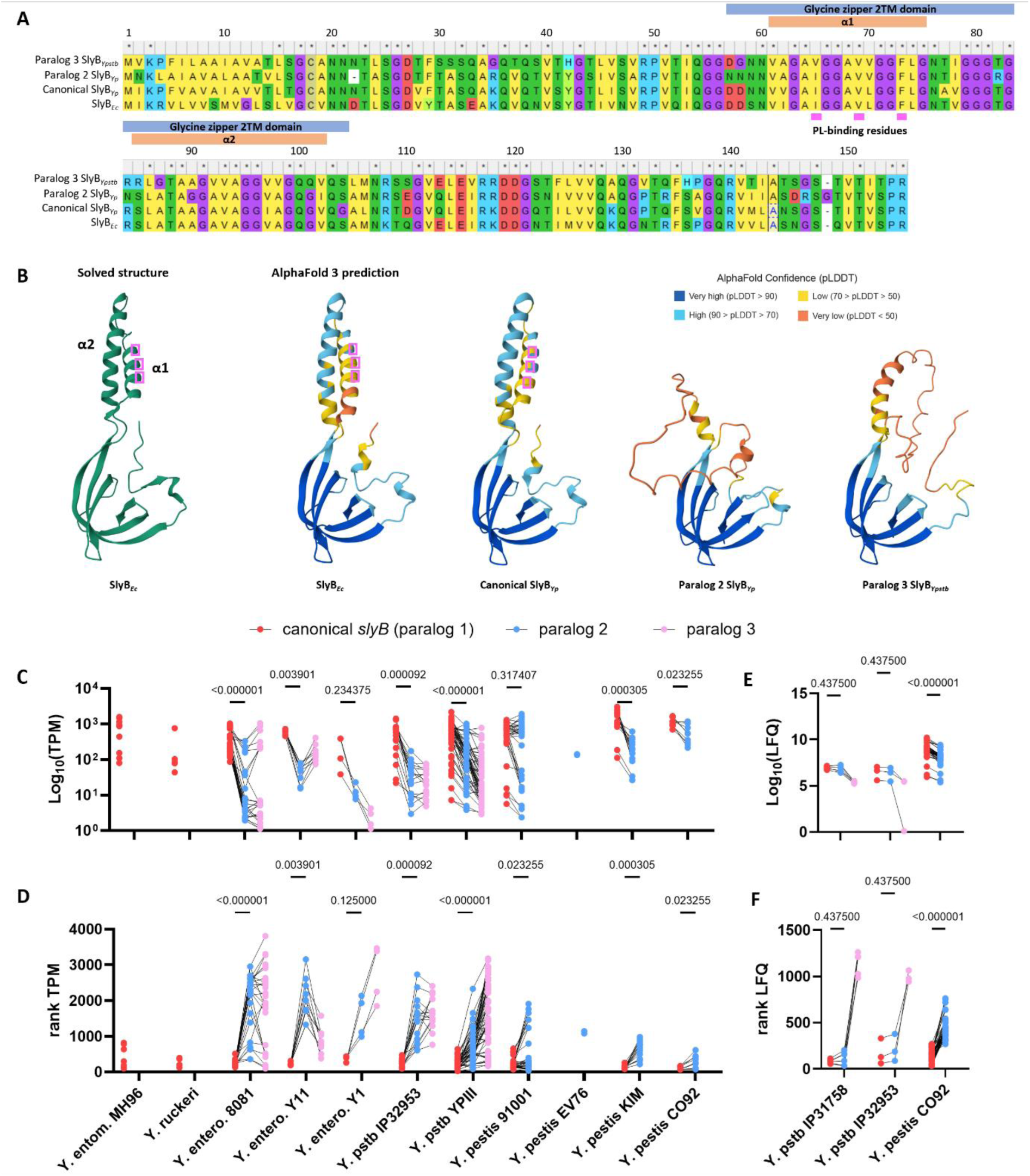
(A) Amino acid alignments and (B) solved and AlphaFold 3 structure prediction of *E. coli* canonical SlyB (SlyB*_Ec_*), *Y. pestis* CO92 canonical SlyB (SlyB*_Yp_*), *Y. pestis* CO92 SlyB paralog 2 and *Y. pseudotuberculosis* IP32593 SlyB (SlyB*_Ypstb_*) paralog 3. Aa numbering is from the 155 aa canonical SlyB*_Ec_* and SlyB*_Yp_*. The glycine zipper 2TM domain and the 2 α-helix containing the glycine zipper are annotated in blue and orange respectively, and the phospholipid (PL)-binding residues are highlighted in pink in both alignments and structures. (C-F) Expression data of the *slyB* paralogs extracted from RNA-Seq-based transcriptomic and LC-MS/MS-based proteomic experiments on *Yersinia* species. Data are plotted as (C) transcripts per million (TPM) and (D) their respective ranking compared to the other transcripts in each experiment for RNA-seq data, and (E) label-free quantification (LFQ) and (F) their respective ranking compared to other proteins in each experiment for LC-MS/MS data. Statistical significance was assessed for expression or rank between canonical *slyB* and the second paralog using the paired non-parametric Wilcoxon test with the with Holm-Šidák correction method for multiple testing. Linked points correspond to transcripts or proteins from the same experiment. Isolate names are the reference genome names to which the data were mapped. Full expression tables can be found in Supplementary Table 5. *Y. ruckeri* data are a pool of SC09 and QMA0440 strain data. entom. : *entomophaga*. entero. : *enterocolitica*. pstb: *pseudotuberculosis*.

Alignment of canonical SlyB orthologs from the *Yersinia* genus also showed conservation of the functional domains during *Yersinia* speciation (Supplementary Fig. 5). Interestingly, the T77A mutation was only present in *Y. pestis* and shared with the most recent – but not with the most ancestral – *Y. pseudotuberculosis* genotypes, as well as some *Y. wautersii* strains. One could hypothesize that the ancestral ACT codon (threonine) mutated to GCT (alanine) in the common ancestor of *Y. similis*, *Y. wautersii* and *Y. pseudotuberculosis*, then independently mutated to GTT (valine) in *Y. similis* and *Y. wautersii* species. A A77T reversion mutation could have occurred during branching with the most ancestral genotype of *Y. pseudotuberculosis*. Based on SlyB*_Ec_* structure, residue 77 is situated between the two α-helix and is linked to the LPS lipid A, suggesting that its mutation could affect binding efficacy to LPS variants.

Altogether, these data show an expansion of SlyB paralogs within the *Yersinia* genus with unclear structural conservation in the paralogs. Concurrently, *Y. pestis* showed paralog 3 inactivation as well as one specific mutation in canonical SlyB emerging during speciation, suggesting a putative adaptation to the pathogen lifestyle.

### Canonical *slyB* is under negative selection in *Y. pestis*

The most recent extensive *Y. pestis* genome datasets from our laboratory and others show that the region encompassing *slyB* and *slyA* (*rovA*) is a hotspot of mutations^46,47^. We further analyzed these datasets encompassing third plague pandemic isolates as well as ancient DNA (aDNA) data from the first and second pandemic to look for selection signals or variants of interest in the two functional *slyB* paralogs. While the mutation hotspot mainly affected *slyA*, the YPO2373 and YPO3646 loci were highly conserved in *Y. pestis* with rare non-synonymous mutations. Ratio of non-synonymous to synonymous substitution rates highlighted negative selection of YPO2373 (dN/dS scores between 0.34 and 0.53) and of YPO3646 to a lower extent (dN/dS scores between 0.68 and 0.95) (Table 1). Two non-synonymous mutations were identified in *Y. pestis* canonical YPO2373 sequence in only 2 strains out of more than 5,000 genomes, at amino acid position 57 and 102 which are conserved residues between SlyB*_Yp_* and SlyB*_Ec_* (Supplementary Table 3 and 4, Fig. 1A). Based on SlyB*_Ec_* structure, D57G mutation could weakly impact the link between the two α-helixes by losing a hydrogen bond with Asp58. The V102I mutation – also present in all *Y. pestis* paralog 2 – is situated in the α2-helix on the OM inner leaflet side, and the carbon addition to the sidechain is unlikely to cause any effect on SlyB function. In the second paralog, YPO3646, only two aa mutations at position A23V and A128S (corresponding to positions 24 and 129 in YPO2373 and SlyB*_Ec_*) were observed in two different isolates. These residues, which may interact, are not conserved between YPO3646, YPO2373 and SlyB*_Ec_* and structural information of SlyB*_Ec_* does not forecast any impact of these mutations. Interestingly, a V92I mutation (position 93 for YPO2373 and SlyB*_Ec_*) was observed in one aDNA sample from the first pandemic^48^. This residue is at the interface between SlyB oligomers, PL and palmitic acid from the anchor lipid and could potentially have an impact on SlyB structure and function. Strikingly, no mutations were observed in the PhoP-binding box and at the transcriptional start site of the canonical locus. Altogether, these results suggest a high selective pressure during *Y. pestis* natural life cycle to preserve canonical SlyB sequence and function across evolution and pandemics.

**Table 1.**
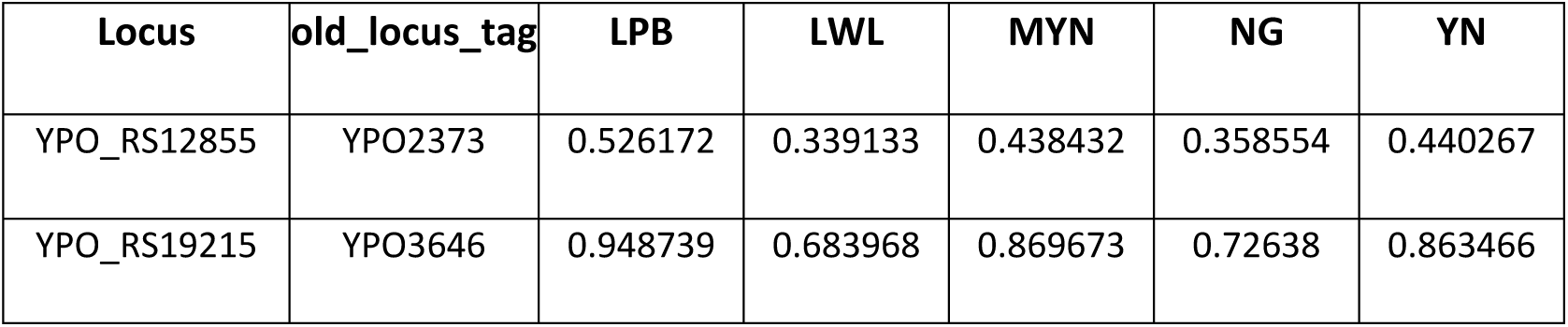
Ratio of non-synonymous to synonymous substitution rates (dN/dS) for canonical *slyB* locus (YPO2373) and paralog 2 (YPO3646) estimated using five different methods.

**Table 2.**
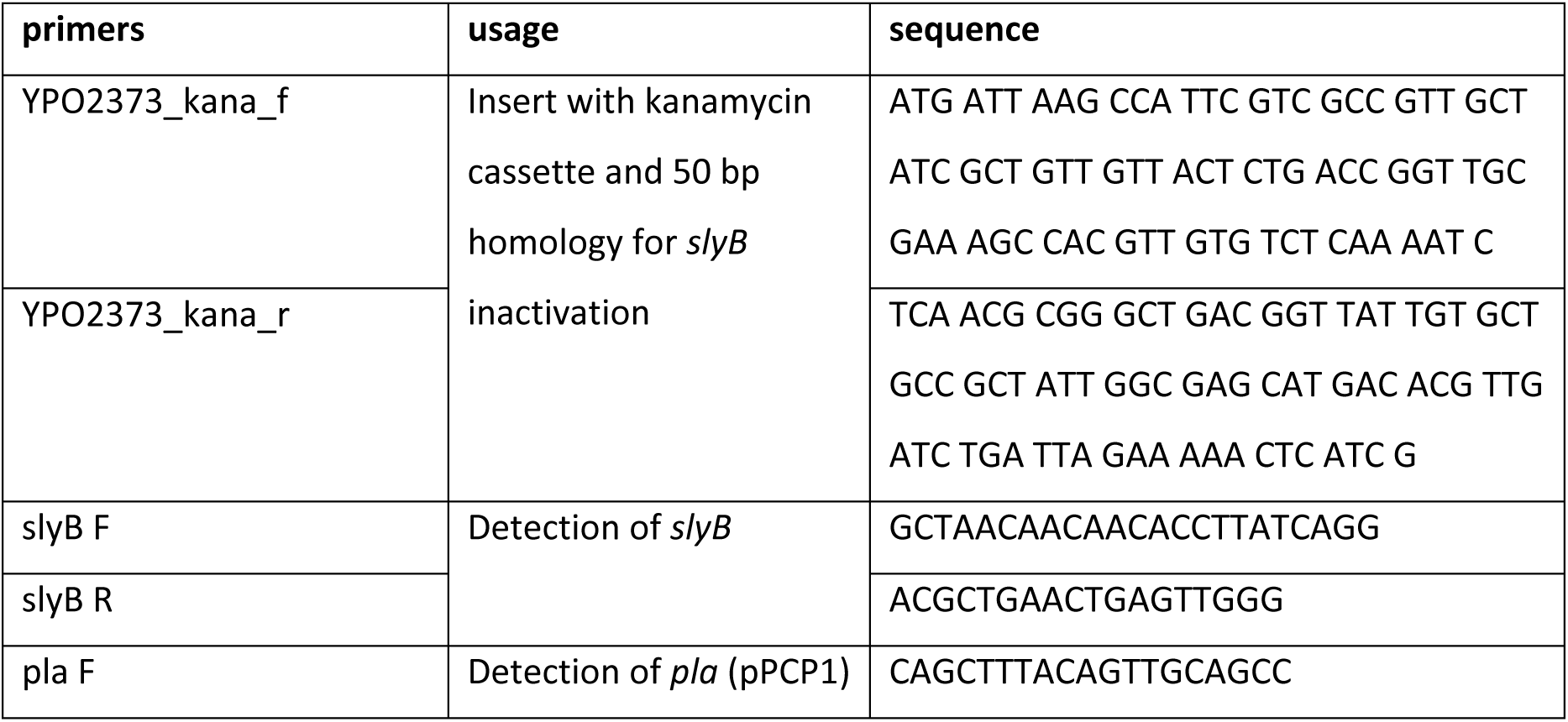
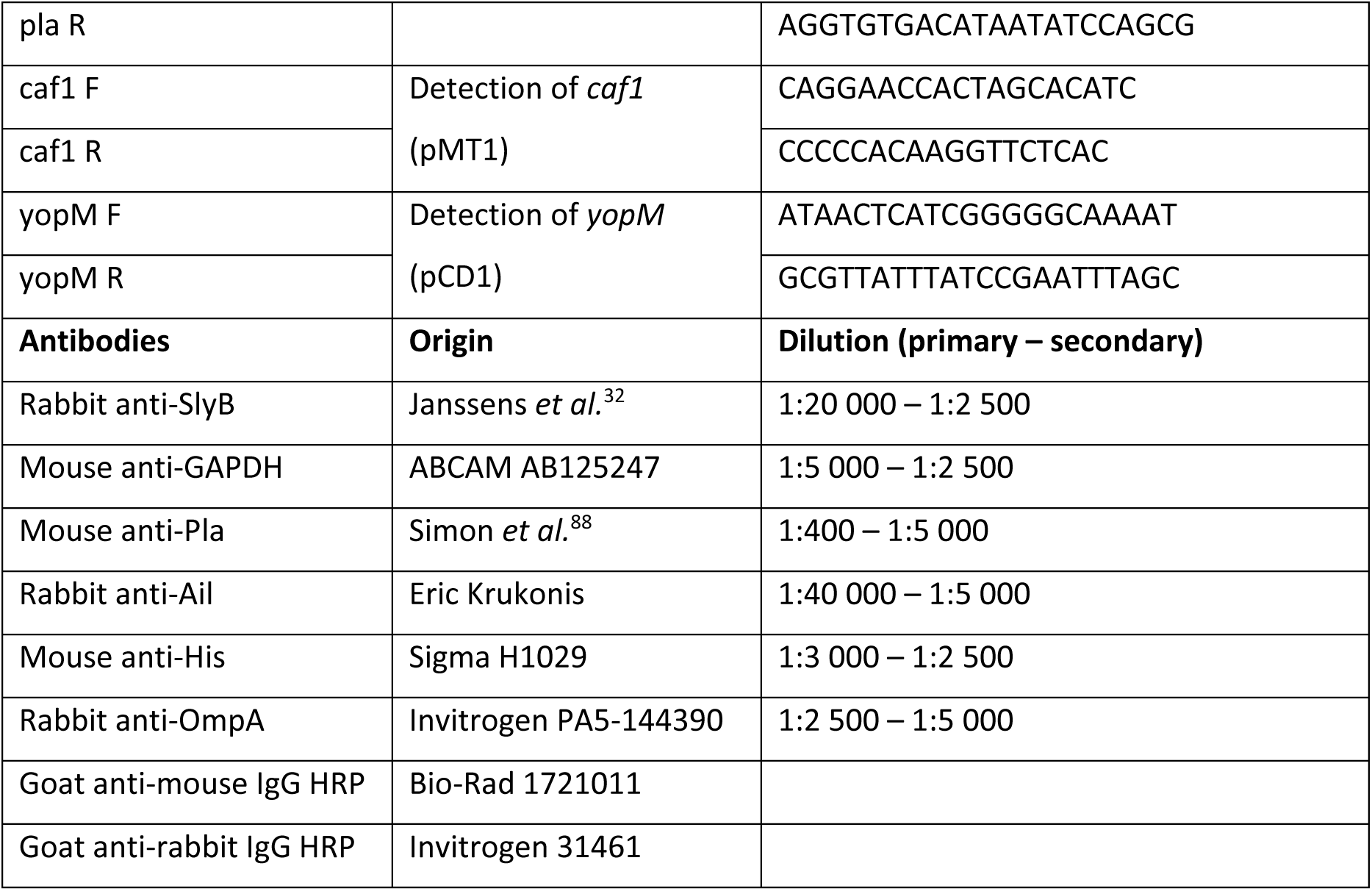
List of primers and antibodies used in this study. HRP: horseradish peroxidase.

### Canonical *slyB* shows high expression patterns across the *Yersinia* genus

To go beyond genomic evolutive analysis and assess paralog comparative usefulness in bacterial lifestyle, we compared the expression of SlyB paralogs across *Yersinia* species based on RNA-sequencing and LC-MS/MS proteomics from the Yersiniomics database^49^ as well as unpublished proteomics data from our laboratory (Fig. 1C-F). These data cover a wide range of species, from the most ancestral *Y. entomophaga* and *Y. ruckeri* to the most recently evolved *Y. pestis*, as well as multiple culture conditions and gene mutations. Overall, the canonical locus was systematically ranked in the first quarter of the most expressed transcripts or proteins, and most of the time in the first 10% most abundant transcripts (ranked below 400 out of around 4000 transcripts), depicting a conserved high requirement of canonical SlyB for *Yersinia* lifestyles. The second and third paralog expression showed lower levels and greater variability in expression. While the second paralog expression was as high or even higher than the one from canonical *slyB* in numerous conditions, especially in *Y. pestis* Microtus (91001) RNA-seq data or *Y. pseudotuberculosis* LC-MS/MS data, both transcriptomics and proteomics data from the more recent *Y. pestis* Orientalis (CO92) or Medievalis (KIM) isolates consistently showed higher expression (approximately 10-fold) of the canonical *slyB* locus and low variability of paralog 2 expression. Interestingly, some laboratory-evolved EV76 vaccinal strains lost the canonical *slyB* locus (as observed in the EV76-CN and EV76-NIIEG complete genomes or formerly by DNA microarray^50^), which was never observed in natural strains. In contrast, the expression patterns of EV76 paralog 2 were consistent with that of paralog 2 from other strains.

Expression data thus point to an important structural role of canonical SlyB in *Yersinia* lifestyle which may be dispensable *in vitro*. Concurrently, SlyB paralogs showing lower basal expression and higher inducibility in *Yersinia* may serve to adapt to specific environmental conditions and stresses. The inducibility seems to be lost in the most recent *Y. pestis* strains, pointing to niche-specific adaptation.

### SlyB function is required for bubonic but not septicemic plague

To investigate the function of the canonical YPO2373 locus, we inactivated the *slyB* gene by allelic exchange using a kanamycin-resistance cassette (Δ*slyB* construct) in *Y. pestis* CO92 strain. We also complemented the mutant strain with a chromosomal insertion of *slyB* and its native promoter (Δ*slyB*::Tn7-*slyB* construct) or with a chromosomal insertion of *slyB* encompassing three point mutations disrupting PL binding in the OM outer leaflet^32^ (I65A, V69A and F73A; Figure 1B), resulting in the Δ*slyB*::Tn7-*slyB*^PL^ construct. These four strains did not show any growth difference in LB at 28°C or 37°C (Supplementary Fig. 6A). SlyB expression levels were similar in CO92 and complemented strains and, as expected, not detected in the Δ*slyB* strain (Supplementary Fig. 6B).

We assessed canonical SlyB requirement along *Y. pestis* lifestyle, starting from mammalian bubonic infection. We observed a robust virulence decrease in two models of bubonic plague in OF1 mice after injection of the Δ*slyB* construct. Indeed, this phenotype was present upon intradermal (ID) or subcutaneous (SC) inoculation (Fig. 2A and B, Supplementary Fig. 7A). Infection with the Δ*slyB*::Tn7-*slyB* construct restored wild-type virulence levels (Fig. 2A and B, Supplementary Fig. 7A). SlyB was recently shown to scaffold lipid nanodomain in the OM, when OM insults result in LPS shedding and flipping of inner leaflet PL to the outer leaflet^32^. To assess if the contribution of SlyB to virulence was linked to this mechanism, mice were infected with the Δ*slyB*::Tn7-*slyB*^PL^ strain described above. This SlyB mutated-strain showed reduced virulence upon both subcutaneous and intradermal inoculation, validating SlyB PL-binding requirement for plague pathogenesis (Fig. 2A and B, Supplementary Fig. 7A). Next, we tested the loss-of-virulence phenotype on a subset of genetically diverse inbred mice from the Collaborative Cross (CC) collection to assess its robustness. Interestingly, the contribution of SlyB to virulence varied from line to line (Supplementary Fig. 7B). More specifically, CC013, CC030, CC042, CC051, CC059 lineages showed similar reduced susceptibility to Δ*slyB* strain infection as OF1 mice. On the other hand, CC061 and CC025 were as susceptible to the Δ*slyB* construct as to WT *Y. pestis* – the latter lineage showing a slight lag in mortality. Finally, CC037 seemed slightly more robust to *Y. pestis* infection overall.

**Figure 2.**
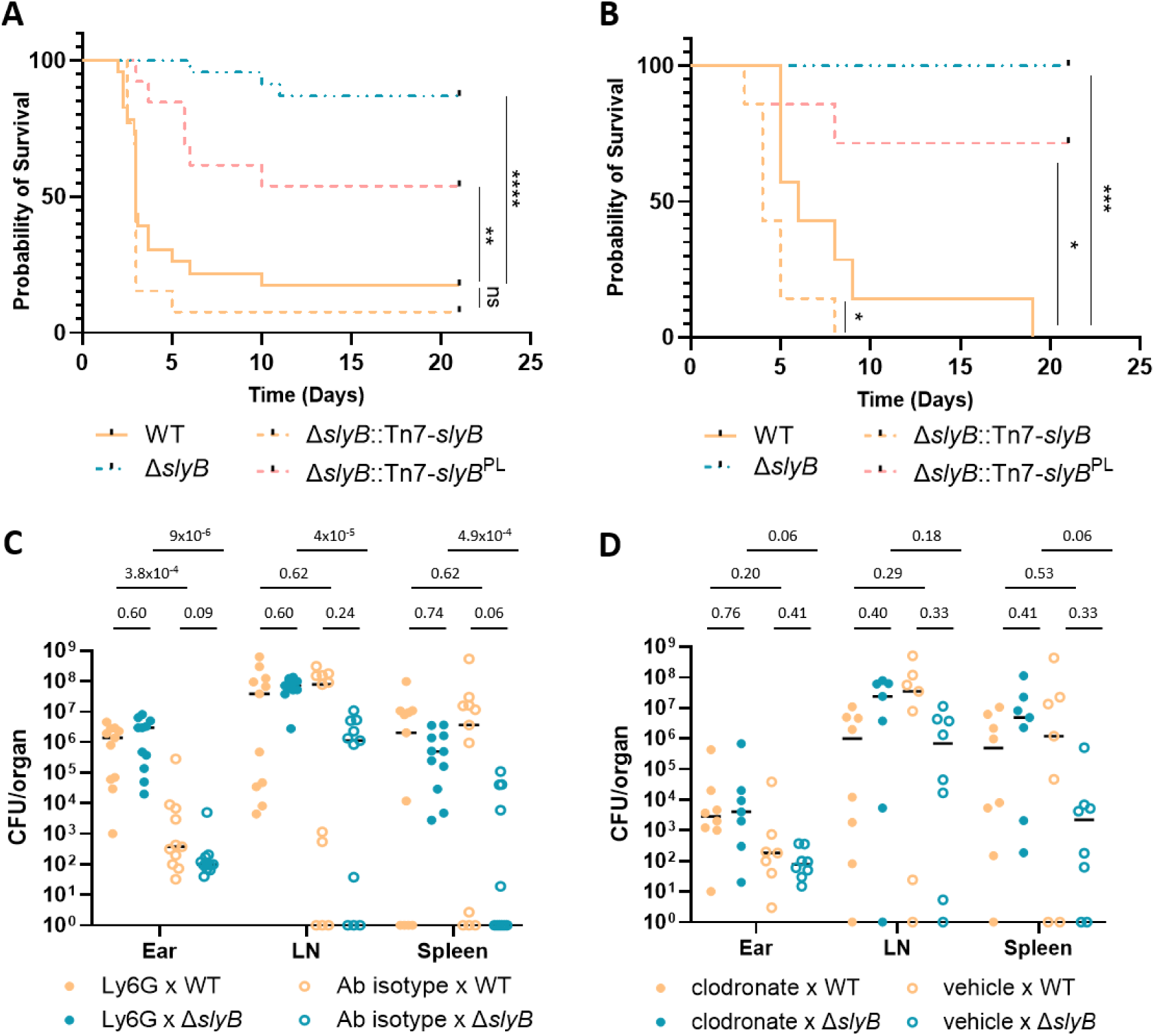
(A) Survival of OF1 mice after intradermal injection in the ear pinna. Data are pooled from 3 to 5 replicate experiments encompassing respectively 13 mice (Δ*slyB*::Tn7-*slyB* and Δ*slyB*::Tn7-*slyB*^PL^) and 23 mice (WT and Δ*slyB*) with injection from 47 to 1,868 CFUs (see Supplementary Fig. 7A for individual experiments). (B) Survival of OF1 mice after subcutaneous injection of approximately 144 to 172 CFUs (one experiment with 7 mice per group). Statistical significance was assessed by Mantel-Cox logrank test. ns: not significant. *: p-value < 0.05. **: p-value < 0.01. ****: p-value < 0.0001. (C) Bacterial load in the ear dermis, draining lymph node (LN) and spleen after neutrophil depletion by intravenous injection of Ly6G-antibody or isotype antibody as control, then intradermal injection in the ear pinna of approximately 225-285 CFUs of the WT and Δ*slyB* construct. Data are pooled from 3 replicate experiments. (D) Bacterial load in the ear dermis, draining lymph node (LN) and spleen after monocyte and macrophage depletion by intravenous and intradermal injection in the ear pinna of clodronate liposome or control liposome (vehicle), then intradermal injection at the same site of approximately 94-313 CFUs of the WT and Δ*slyB* construct. Data are pooled from 2 replicate experiments. A non-parametric multiple Mann-Whitney test with Holm-Šidák correction method was performed for each pairwise comparison on the three organs.

It was previously shown that a strong dissemination bottleneck occurs in the dermis upon ID injection of *Y. pestis*, while this was bypassed upon SC inoculation^51^. Since SlyB contributed to virulence upon both ID and SC injection, we hypothesized that SlyB was not required for escape from the dermis but at a later stage of infection, either at the lymph node levels or during hematogenous dissemination. We estimated the bacterial load in the dLN and in the spleen after intradermal injection in the ear pinna and observed a reduced amount of SlyB mutant in both organs (Supplementary Fig. 7C). Concurrently, we did not observe any difference in virulence between the WT and mutant strain after intravenous injection (Supplementary Fig. 8A). The mouse complement system being less potent to kill *Y. pestis* than other mammalian blood^24^, we also tested Δ*slyB* strain capacity to survive in human blood. The Δ*slyB* strain could grow in normal human serum and inactivated human serum at the same rate as the WT strain at 37°C, despite a slight survival decrease at the late stationary phase which was not observed when grown in LB supplemented with calcium at 37°C (Supplementary Fig. 8B and C). We could not reproducibly observe differences for survival in human whole blood from different donors with two different anticoagulants (Supplementary Fig. 8D). These data suggest that *Y. pestis* SlyB requirement is restricted to the dLN colonization and neither dermis egress nor survival and replication into the blood.

### SlyB is required for mammalian but not to arthropod immune system resistance

We next sought to identify from which key immune cells SlyB could allow to escape. We depleted neutrophils or monocytes in mice and assessed bacterial colonization of Δ*slyB* in the dermis at the injection point, in the dLN and in the spleen. Antibody-mediated neutrophil depletion showed an increased bacterial replication for both WT and Δ*slyB* strain in the dermis 48 hours post-infection, confirming former results showing that neutrophils restricted bacterial replication at the injection point^5^. In the dLN but not in the spleen, Δ*slyB* replication was restored to WT levels after neutrophil depletion, suggesting SlyB is necessary to resist neutrophils antimicrobial activity in the dLN (Fig. 2C). Local macrophages and circulating monocyte depletion by liposome clodronate injections led to increased bacterial load of the Δ*slyB* strain in the spleen and in the dLN, although these data were more variable, suggesting a role of monocytes in Δ*slyB* replication control (Fig. 2D). To get deeper insights in specific immune stresses preventing Δ*slyB* strain colonization, we tested intracellular and extracellular survival of bacteria following *in vitro* exposure to human macrophages and neutrophils, RAW 264.7 macrophages, or isolated effector molecules such as neutrophil compounds (neutrophil extracellular traps, granules, the cathelicidin LL-37) or H_2_O_2_. However, unexpectedly, we could not detect any difference between the WT and Δ*slyB* strains (Supplementary Fig. 9). This suggest that SlyB could protect bacteria from other antibacterial compounds or compound combinations, or that it is required in the more complex *in vivo* environment during lymph node colonization, considering the kinetics of environmental changes – such as temperature switch – as well as from the immune response.

To cover the whole ecological cycle of *Y. pestis*, we assessed canonical SlyB requirement for flea colonization and biofilm formation at environmental temperature. The Δ*slyB* strain was perfectly able to colonize the flea proventriculus, showing a morphology and level of colonization comparable to the WT strain (Fig. 3A and B). Flea competitive infection with both strains did not show any advantages for the WT on the mutant strain, even showing a very slight but significant increase in Δ*slyB* mutant colonization after 13 days of infection compared to day 0 (Fig. 3C). Concurrently, the Δ*slyB* construct had a higher capacity to form biofilm *in vitro* at environmental temperature (Fig. 3D). As biofilm formation is regulated by c-di-GMP in *Y. pestis*, we quantified intracellular c-di-GMP *in vitro* in the Δ*slyB* and Δ*slyB*::Tn7-*slyB* constructs but could not observe any differences (Fig. 3E), pointing to other regulating pathways.

**Figure 3.**
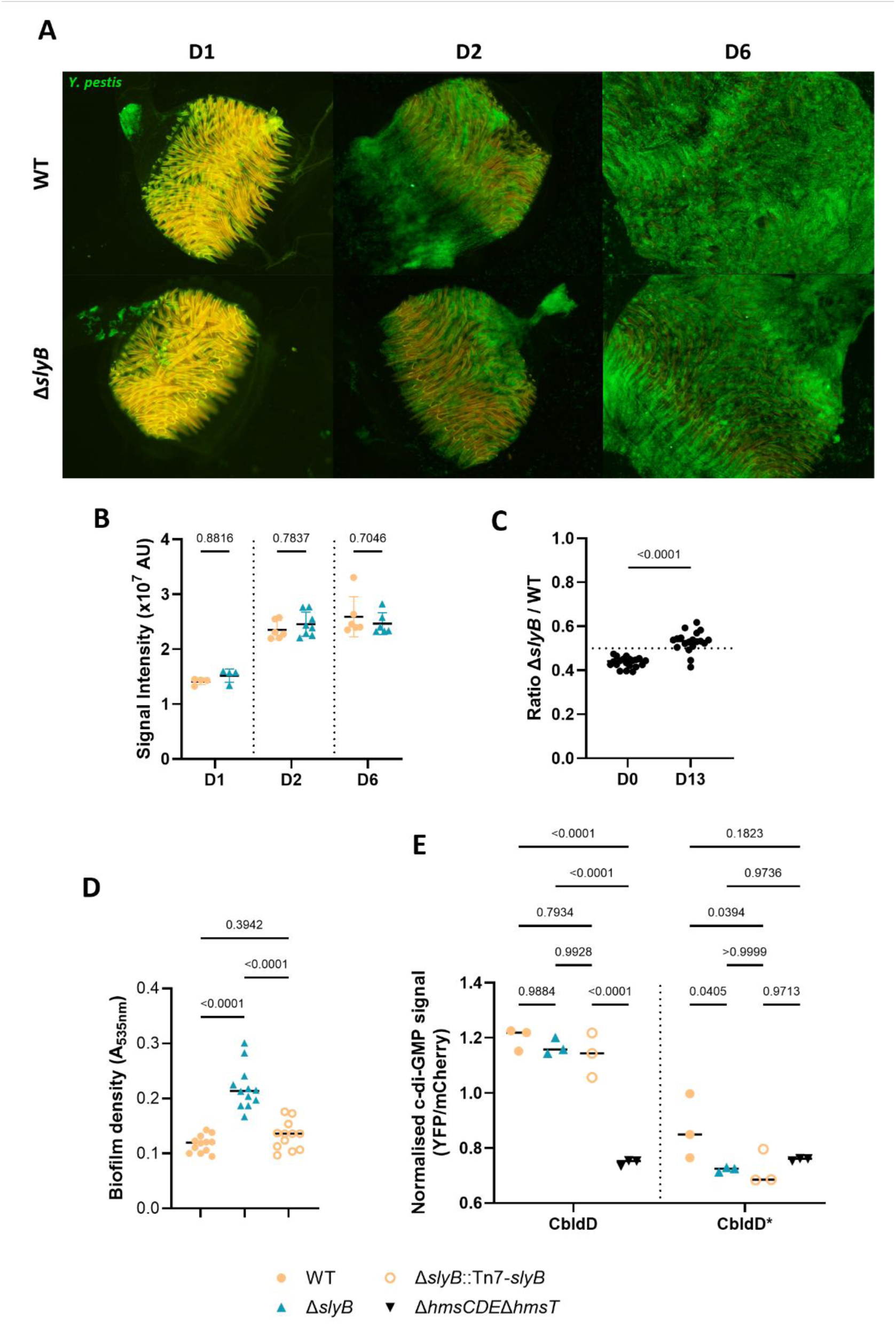
(A) Representative pictures of flea proventriculus colonization by WT and Δ*slyB Y. pestis*-GFP at days 1, 2 and 6 post flea feeding and (B) its associated quantification. Statistics were computed by two-way ANOVA with Šidák’s multiple comparisons test. (C) Ratio of Δ*slyB* construct on WT strains during flea competitive infection with the same amount of both strains at days 0 and 13 post flea feeding. Statistics were computed by non-parametric Mann-Whitney test. (D) Biofilm density after 48h growth of WT, Δ*slyB* and Δ*slyB*::Tn7-*slyB* constructs at 21°C, assessed by crystal violet staining. Statistics were computed by one-way ANOVA with Tukey’s multiple comparisons test. (E) Intracellular c-di-GMP in *in vitro* biofilm, measured by YFP fluorescence after c-di-GMP-dependent dimerization of C-terminal domain of BldD (CBldD), normalized by mCherry fluorescence. CBldD* is unable to bind c-di-GMP and to dimerize, while Δ*hmsCDE*Δ*hmsT* strain is unable to produce c-di-GMP. Statistics were computed by two-way ANOVA with Šidák’s multiple comparisons test.

SlyB thus seems to be required for immune system escape during mammalian host colonization at 37°C, whereas its absence does not preclude flea proventriculus colonization at environmental temperature.

### Canonical SlyB function is specifically required for polymyxin B resistance at mammalian but not environmental temperature

Previous experiments on SlyB mutants in other bacteria showed reduced resistance to antimicrobial peptides (AMP) such as polymyxin B (PmB), cathelicidin LL-37 or bactenecin-derived Bac2A^32,41^. AMP are amphipathic molecules destabilizing the outer membrane of diderm bacteria and serving as the most ancestral immune system in both arthropods and mammalians. *Y. pestis* is naturally resistant to PmB at environmental temperatures mainly due to LPS modifications (minimal inhibitory concentration – MIC >128 mg/L), but it shows a marked increase in susceptibility at mammalian temperature (MIC around 1 or 2 mg/L). While we could not detect any difference in LL-37 susceptibility for *Y. pestis* Δ*slyB* construct (Supplementary Fig. 9D), we observed a slight increase in PmB susceptibility when exposed at 37°C but not at 28°C (Fig. 4A). This was the case for both bacteria precultivated at 28°C and 37°C before exposition to PmB at 28°C or 37°C. This phenotype was not dependent on the presence of the pYV- plasmid (harboring the T3SS) but was dependent on the 3 PL-binding residues, as shown with the Δ*slyB*::Tn7-*slyB*^PL^ construct. Using disk diffusion assay at 28°C and 37°C, we also assessed Δ*slyB* construct susceptibility to PmB and 15 other antibiotics from 13 different classes and subclasses targeting peptidoglycan metabolism, gyrase, folate pathway, or RNA and protein synthesis. No clear difference in susceptibility to any antibiotics could be observed at both temperatures, suggesting SlyB is not involved in resistance to these other types of cellular stress. Unlike with PmB, we could not observe any difference in polymyxin E (colistin) sensitivity, which may be due to poor cAMP diffusion and to *Y. pestis* natural resistance to this antibiotic, decreasing colistin diffusion assay resolution, or to the slight sequence difference between PmB and colistin (Supplementary Fig. 11).

**Figure 4.**
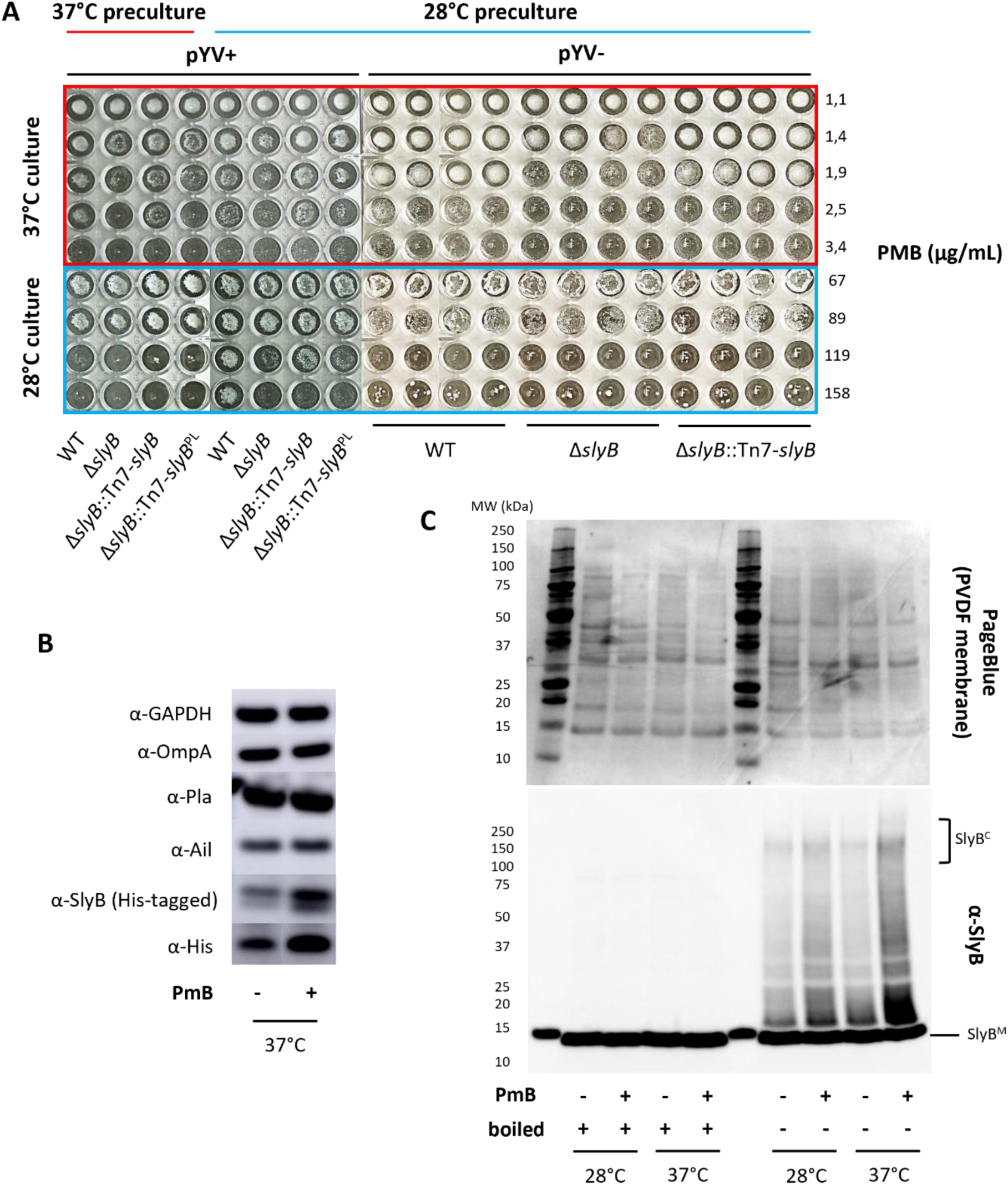
(A) Polymyxin B (PmB) minimum inhibitory concentration (MIC) for diverse *slyB* constructs. After preculture at mid-exponential phase in LB at 28°C or 37°C, PmB exposition was done for 48h at 28°C and 37°C in cation-adjusted Mueller-Hinton (MH II) media. Full pictures and conditions are shown in Supplementary Fig 10. (B) Western blot (composite figure) against GAPDH, diverse abundant outer membrane proteins (OmpA, Pla, Ail), SlyB and His-tag. His-tagged SlyB strain (cured from pYV plasmid) was exposed or not to 2 µg/mL PmB for 1 hour at 37°C in MH II media under agitation. Anti-His western blot was performed independently of the other western blots. (C) Semi-native polyacrylamide gel electrophoresis (snPAGE) transferred on PVDF membrane and colored with PageBlue, and its associated anti-SlyB western blot on bacterial whole cell lysate. CO92 strain (cured from pYV plasmid) was exposed or not to 10 µg/mL PmB for 1 hour at 28°C or 37°C in MH II media under agitation. MW: molecular weight. SlyB^C^: SlyB complexes. SlyB^M^: SlyB monomers.

Transient exposure to PmB at 37°C led to an increased abundance of SlyB, while the most abundant *Y. pestis* OMP such as OmpA, Pla or Ail did not show any abundance changes by western blotting (Fig. 4B). This is expected as canonical SlyB is regulated by the PhoPQ TCS^37^, which may sense AMP^52^. As we observed a second band which could stem from a SlyB paralog, we complemented the Δ*slyB* strain with a His-tagged canonical SlyB (Δ*slyB*::Tn7-*slyB*-TEV-His construct), which did not change PmB susceptibility (Supplementary Fig. 10), and could thus validate induction of the canonical SlyB (Fig. 4B). By performing semi-native polyacrylamide gel electrophoresis (snPAGE), we could observe the formation of high molecular weight SlyB complexes upon PmB exposure, recapitulating SlyB lipid nanodomain formation induced by other membrane stresses in *E. coli*^32^ (Fig. 4C). This induction was present at both 28°C and 37°C but was higher at the latter temperature, confirming SlyB function to be most important during mammalian infection.

## Discussion

SlyB was described as a mediator of OM homeostasis across the Proteobacteria phylum^32,41,45^ and in the core-regulon of the PhoPQ TCS^37^. Genetic linkage of *slyB* and the MarR-family transcription factor (TF) encoding gene *slyA* (also named *rovA*) was additionally described in most species of the *Enterobacterales* order^36^. Within the *Yersinia* genus, the ancestral state – a single canonical *slyB* allele situated adjacent to *slyA* – was maintained in early-branching species such as *Y. entomophaga* and *Y. ruckeri*. After their divergence, environmentally versatile lineages acquired two additional paralogs inserted near loci involved in cold adaptation (*cspAG*), LPS remodeling (*lpxP*), and motility (*mot*). These paralogs showed unclear structural conservation, and exhibit lower basal expression and greater condition-dependent variability than the PhoPQ-regulated canonical copy. This suggests that these paralogs could assume other functions, or they could serve to fine-tune SlyB dosage across fluctuating environments such as variation in temperature, salinity or pH, varying host or biosphere colonization with varying microbial competition. Why this expansion specifically happened in the *Yersinia* genus, whether paralogs retain identical nanodomain-scaffolding functions or have assumed divergent structural roles, and to which membrane stresses these paralogs could respond in the former case, remain to be determined.

After the transition from the enteric and environmental lifestyle of *Y. pseudotuberculosis* to the flea-rodent cycle of *Y. pestis*, the third paralog was inactivated by frameshift mutation and the second paralog abundance was consistently reduced, as shown by shotgun proteomics data. In contrast, the canonical locus remained under strong purifying selection in *Y. pestis* (dN/dS < 0.6). This dichotomy between paralogs mirrors the well-known genome-wide trajectory of *Y. pesti*s niche specialization, leading to loss of metabolic flexibility coupled with conservation of structural and virulence determinants^53^. The loss of the entire *slyA/slyB* region in laboratory-adapted EV76 vaccine strains^50^ – never documented in natural isolates – further underscores the ecological requirement of this locus despite being dispensable *in vitro*. However, the relative contributions of *slyB* versus *slyA* to fitness in the wild remain unresolved, the latter being a well-known TF involved in *Yersinia* virulence^54–56^. Concurrently, serial *in vitro* passage at 28°C leading to downregulation or loss of the abundant OMP virulence factors Ail and Pla correlates with overexpression of OmpA, OmpF, OmpC or SlyB, suggesting that compensatory envelope modulation is also engaged during laboratory adaptation^57^. Experimental *slyA* deletion additionally led to dysregulation of envelop-related genes and upregulation of *slyB* and T3SS-encoding genes at 37°C, suggesting an indirect role of *slyA* in envelope homeostasis^58^. Interestingly, recent reports from our laboratory and others additionally highlighted a mutation hotspot in the *slyA/slyB* region of *Y. pestis*^46,47^. SlyA probably evolved under strong positive selection with concomitant regulon rewiring^36^, whereas we eventually showed here that SlyB was under negative selection as a structural protein for which sequence fidelity was strongly enforced. Altogether, these observations highlight the importance of the *slyA/slyB* locus in *Y. pestis* ecology, and we further explored SlyB role in plague natural history.

In the flea vector, the Δ*slyB* mutant exhibited wild-type proventricular colonization and competitive fitness. *Y. pestis* is known to resist the antimicrobial peptide-based arthropod immune system primarily through PhoPQ-dependent addition of Ara4N to hexa-acylated lipid A at environmental temperature^14,15^, a mechanism probably sufficient to prevent envelope destabilization without SlyB-mediated lipid nanodomain assembly. Interestingly, the mutant displayed enhanced biofilm formation *in vitro* at 21°C. A direct polar effect of *slyB* mutation due to the proximity with *slyA*, shown to represses *hmsT* and biofilm production^59^, was excluded as the phenotype was independent of the canonical c-di-GMP pathway and as the function of *slyB* could be restored by expression in *trans* in the Δ*slyB* strain. The lack of SlyB could activate alternative signaling cascades that promote matrix production, but the exact mechanism of SlyB-dependent biofilm inhibition remains to be elucidated.

The requirement for SlyB emerges specifically during mammalian infection at 37°C. Upon host entry, *Y. pestis* shifts to tetra-acylated LPS through loss of PagP and LpxL activity, a remodeling strategy that attenuates TLR4 recognition^16–18^, but could increase intrinsic OM fluidity and permeability^25,60^. PagP deficiency – abrogating palmitate transfer from PL donor to lipid A – as well as reduced expression of phospholipase A PldA at 37°C may further elevate PL content in the outer leaflet, promoting OM symmetrization and compromising barrier function during membrane stress^25^. We propose that SlyB counterbalances these structural liabilities by scaffolding stress-responsive lipid nanodomains, as evidenced by its requirement for PmB resistance and PmB-induced oligomerization at 37°C – a process dependent on its three PL-binding residues and recapitulated by the intermediate virulence defect of the PL-binding-deficient mutant. Thus, *Y. pestis* LPS evolution toward mammalian immune escape appears to have imposed additional pressure on SlyB-dependent envelope resilience. A T77A substitution at the LPS-binding interface, shared by *Y. pestis* and its most recent *Y. pseudotuberculosis* ancestors, could reflect adaptive tuning for the tetra-acylated lipid A, or alternatively cooperative interactions with abundant OM proteins important in virulence and/or envelop structural integrity such as Ail^25,26,61^ or Pla^23^.

Within the draining lymph node, *Y. pestis* expresses the T3SS and Pla to subvert phagocyte function and replicate^5,6,62^, yet massive neutrophil recruitment may generate a hostile microenvironment rich in antimicrobial compounds and oxidative stress. Prior studies established that PhoPQ-dependent resistance to neutrophil granules is only partially attributable to Ara4N decoration or other defined effectors^19,63,64^. Our data identify SlyB as an additional PhoPQ-regulated factor required to withstand neutrophil-mediated killing during bubonic plague, as neutrophil depletion restored Δ*slyB* replication to wild-type levels in the lymph node. The exact molecular effectors remain elusive, as no single *in vitro* stress – including H_2_O_2_, LL-37, or phagocyte infection – recapitulated the defect. This implies that SlyB is necessary to resist another unidentified stress factor, and/or specifically during *in vivo* infection kinetics and combinatorial stresses in tissues, integrating the simultaneous environment and temperature transition inducing LPS remodeling, as well as recruitment dynamics of immune cells. We cannot exclude, however, that restored bacterial burdens following neutrophil depletion partly reflect unchecked dermal replication and continuous draining from injection point to the dLN, rather than intrinsic lymph-node-specific rescue alone.

Surprisingly, SlyB is not required for the bacteremic phase. The Δ*slyB* mutant survived intravenous challenge and replicated in human whole blood and serum at 37°C. Although neutrophils and monocytes should remain bactericidal in blood, high concentrations of glucose and divalent cations in plasma may independently stabilize membrane integrity, as previously shown with a Δ*ail* mutant^25^. Moreover, complement is efficiently neutralized by Ail and SubB^26,65^. The absence of the acute, multifactorial envelope stress likely happening in the lymph node could render SlyB scaffolding redundant during high-titer bacteremia. This underscores the niche-specific requirement of virulence factors across the plague cycle.

The contribution of SlyB is further nuanced by host genetics, as Collaborative Cross mice displayed variable susceptibility to Δ*slyB* infection. This variability in susceptibility could stem from specific unidentified mutation or alleles in gene(s) involved in bacterial control as we previously described with *Y. pestis*^66,67^, and as described elsewhere with other infections (CD11a-encoding gene *Itgal* in CC042 lineage^68,69^, Ncf2, Slc11a1^70^, and others^71–73^). Alternatively, differences in specific immune cell basal level and/or infection-mediated induction could explain why some CC strains are unable to control Δ*slyB* strain replication^74,75^. Future immunophenotyping of our CC collection and expression quantitative trait locus mapping should pinpoint host determinants that impose or alleviate SlyB-dependent envelope stress. The weak but persistent expression of the second paralog YPO3646 also raises the possibility of partially redundant functions that merit investigation. Together with Ail, which stabilizes LPS at 37°C^25,61^, canonical SlyB thus appears to sustain the delicate equilibrium between immune evasion and structural integrity that defines the *Y. pestis* OM.

In conclusion, canonical SlyB may have been evolutionarily refined during *Y. pestis* emergence to meet a precise ecological demand: resisting immunity-driven envelope stress during dissemination in the mammalian host. Its dispensability in the flea vector and during septicemia emphasizes that even structural factors can evolve to be deployed in a stage-specific manner. SlyB thus emerges as a broadly conserved structural protein in environmental and pathogenic bacteria, which function could be coopted and fine-tuned to varying lifestyles inside and across species. These findings may also integrate SlyB into the PhoPQ-regulated envelope defense network and establish lipid nanodomain encapsulation as a critical axis of bubonic plague pathogenesis.

## Material and Methods

### Phylogenetic and structure analysis

Phylogenetic reconstruction was performed using concatenated nucleotide sequences of the 500 genes of the cgMLST *Yersinia*^76^. The analysis included 374 isolates representing the 28 species of the *Yersinia* genus^77^, including the very recently described *Y. fenwicki*^78^ and *Y. antarctica* (in revision). Concatenated sequences were aligned using MAFFT v7.467 and a maximum-likelihood phylogeny was inferred using IQ-TREE v2.4.0, with ModelFinder to estimate the best substitution model^79,80^.

The presence of the *slyB* gene and its two paralogs was screened across the 374 genomes using ABRicate v 1.2.0 (https://github.com/tseemann/abricate). A custom database was constructed containing the following reference sequences from the *Y. pestis* CO92 reference genome (accession number NC_003143.1): *slyB* (locus tag YPO_RS12855), paralog2 (locus tag YPO_RS19215), and paralog3 (locus tag YPO_RS04730). Hits were retained based on thresholds of ≥ 60 % nucleotide identity and ≥ 60 % query coverage. The phylogenetic tree was visualized and annotated with gene presence data using iTOL v7^81^. Amino acid alignments were realized with MEGA v12 software^82^.

Point mutations in canonical *slyB* and on paralog 2 were identified across the genomes of 5,744 isolates obtained from Mas Fiol *et al.*^46^, using RedDog pipeline (https://github.com/scwatts/reddog-nf) by mapping the sequences onto *Y. pestis* CO92 reference genome (accession number NC_003143.1). KaKs_Calculator v3.0^83^ was then used to calculate dN/dS ratios for canonical *slyB* and paralog 2 using all five models (GLWL, GMYN, GNG, GLPB and GYN). The status of paralog 3 was assessed using blast searches in ABRicate v1.2.0 on the genomic assemblies, and then plotted onto the phylogenetic tree of the isolates with Treeio v1.28.0 R-package^84^.

AlphaFold predictions were realized using AlphaFold server powered by AlphaFold 3 and after removal of the signal peptide. SlyB*_Ec_*solved structure was retrieved from the Protein Database (PDB 7OJG).

### Bacterial strains, culture media and blood

The virulent *Y. pestis* CO92 strain was used in this study^85^. Bacteria were routinely grown on LB agar plates supplemented with 0.002% pork hemin (LBH) at 25-28°C if not stated otherwise. Frozen human serum and buffy coats were provided by the Etablissement Français du Sang (EFS), and fresh human blood (anticoagulated with heparin or ACD) were provided by EFS or the Clinical Investigation Center INVOLvE (Investigation and volunteers for human health) at Institut Pasteur^86^. At INVOLvE, human peripheral blood samples were collected from healthy volunteers. The participants received oral and written information about the research and gave written informed consent in the frame of the COSIPOP cohort after approval of the CPP Est II Ethics Committee (2023, Feb 20^th^). For bacterial enumeration, bacteria were serially diluted 1/10 (20 μl in 180 μL) in phosphate-buffered saline (PBS) and plated on LBH agar plate. Colony-forming units (CFU) were enumerated after incubation at 28°C for 48 hours, or at room temperature for 3 to 4 days. Strains were curated from the pYV plasmid after overnight growth in LB at 37°C followed by plating on magnesium oxalate (MOX) agar plate and selection of colonies growing at 37°C, then verified by PCR and mice survival after subcutaneous injection of 10^6^ CFUs.

### Mutagenesis and complementation

Gene deletion was performed using homologous recombination system^87^. Forward oligonucleotides were designed with around 50 nucleotides of homology with the beginning of the target genes and around 20 nucleotides of the beginning a kanamycin resistance cassette. The reverse oligonucleotides were designed with around 50 nucleotides of homology with the end of the target genes and around 20 nucleotides of the end of the kanamycin resistance cassette. PCR products were precipitated overnight in potassium acetate and ethanol at −20°C, centrifugated at 15,000 g for 30 minutes at 4°C, then 1 mL of 70% ethanol was added and centrifugated at 15,000 g for 15 minutes at 4°C. Supernatant was discarded and DNA pellets were dried for 15 minutes in a SpeedVac. The DNA pellets were resuspended in 5 μL of milliQ water and dialyzed on 0.075 μm filter for 30 minutes. Dyalized DNAs were electroporated in electrocompetent *Y. pestis* CO92 strain harboring the pKOBEG-*sacB* plasmid. Recombinant clones harboring the resistance cassette were selected on LBH plates containing 30 μg/mL kanamycin. Absence of the targeted genes was assessed by PCR using primers inside the target, and insertion of the cassette was assessed by PCR on the flanking region, using primers outside of the target and inside of the cassette. Curation of the pKOBEG-*sacB* plasmid was performed by growing clones on LBH plate without sodium chloride and with 10% of sucrose. Presence of the virulence plasmids was confirmed by PCR on the *yopM* (pCD1), *caf* (pMT1) and *pla* (pPCP1) genes. The *slyB* mutant was complemented in *trans* by cloning the gene (or various constructs) and its upstream and downstream regions in the pUC18R6K plasmid, which does not replicate in *Y. pestis* and harbors the miniTn7 transposon with a chloramphenicol resistance cassette. Electrocompetent cells of the Δ*slyB* strain were electroporated with 400 ng of pUC18R6K-miniTn7::*slyB* (or various constructs) and 400 ng of pTNS2 harboring the transposase and which does not replicate in *Y. pestis*. Selection for the insertion of the miniTn7 was done on LBH plates containing 25 μg/mL chloramphenicol. Verification of the presence of *slyB* was done by PCR with primers inside *slyB*, and insertion of the miniTn7 was verified with PCR on the flanking regions. Absence of the pUC18R6K-miniTn7::*slyB* and of the pTNS2 was assessed by streaking bacteria on LBH with 100 μg/mL carbenicillin and PCR on the *bla* resistance cassette. Strain genomes were analyzed by Nanopore MinION sequencing using SQK-RBK114.24 kit and R10 flow cell to validate the constructs. For flea experiments, mutants were reconstructed in a CO92 pYV- background as explained above following the protocol described in Bontemps-Gallo *et al.*^21^. PCR products were purified using NucleoSpin Gel and PCR Clean-up (Macherey-Nagel) following manufacturer instruction. DNAs were electroporated in electrocompetent *Y. pestis* CO92 pYV- strain harboring the pEP1436 plasmid. Recombinant clones harboring the resistance cassette were selected on LB plates containing 30 μg/mL Trimethoprim. Absence of the targeted genes was assessed by PCR using primers inside the target, and insertion of the cassette was assessed by PCR on the flanking region, using primers outside of the target and inside of the cassette. Curation of the pEP1436 plasmid was performed by growing clones on LB plate with 10% of sucrose.

### Infection of mice

Female OF1 mice (7-week-old) were purchased from Charles River Laboratory (L’Arbresle, France) and Collaborative Cross mice were acquired from Institut Pasteur breeding facility. Prior to infection, they were maintained in the Institut Pasteur BSL-3 animal facility, accredited by the French Ministry of Agriculture (B 75 15-01, issued on May 22, 2008), in compliance with French and European regulations (EC Directive 86/609, French Law 2001-486; June 6, 2001). The research protocol was approved by the French Ministry of Research (N° 2023-0104). Bacterial glycerol stock was thawed from −80°C in LB and grown overnight at 28°C under agitation, then plated and incubated for 24 hours on LBH plates at 25-28°C. Bacteria were resuspended in 3 mL PBS, vortexed for 1 minute and the optical density at 600 nm (OD_600_) was adjusted in 10 mL PBS and serially diluted to reach the desired concentration. For intravenous injections, mouse tails were heated with a paper soaked in warm water and 100 μL of bacterial suspension were injected in the caudal vein with a 29G needle. For intradermal injection, mice were anesthetized by intraperitoneal or subcutaneous injection of 100 µL xylazine (10 mg/kg) and ketamine (100 mg/kg) and 2 μL of the bacterial suspension were injected in the ear pinna with a NanoFil™ syringe (World Precision Instrument). The inoculum was verified by plating dilutions on LBH plates and incubation at 28°C for 48 hours. Survival of the mice was monitored for a minimum of 17 days.

Depletion of neutrophils was realized by intravenous injection of 250 µg Ly6G/Ly6C antibody (*InVivo*MAb NIMP-R14 clone) or isotype control (Rat IgG2b) one day before and one day after infection. Efficient depletion was verified by cytometry one day after injection in uninfected mice. Depletion of circulating monocyte and resident macrophage was realized by simultaneous intravenous and intradermal injection of respectively 10 µg (in 2 µL) or 40 mg/kg clodronate liposome or empty liposome control (Encapsula NanoSciences Clodrosome® + Encapsome®) 2 days prior to infection. Organs were harvested shortly after mouse euthanasia with CO_2_. Ears were cut, and the two dermis sheets were separated and homogenized by bead beating with a Precellys 24 tissue homogenizer (Bertin technologies) in 500 µL PBS. Superficial parotid lymph node and spleen were harvested in GentleMACS® M Tube filled with 2 mL PBS then homogenized in an Octo Dissociator (Miltenyi Biotec, Bergisch Gladbach, Germany) using RNA_2 program. Homogenate were serial diluted, plated on LBH (with or without supplementation with 1 µg/mL irgasan) and incubated for 2 days at 28°C or 3 days at 21°C.

### Survival in human blood and serum

Parental CO92 and mutant strain glycerol stocks were thawed from −80°C in LB and grown overnight at 28°C under agitation, then plated and incubated for 24 hours on LBH plates at 25-28°C. Bacteria were resuspended in 3 mL PBS, vortexed for 1 minute and the optical density at 600nm (OD_600_) was adjusted to 0.25 in 10 mL PBS to reach a concentration around 2×10^8^ CFU/mL. Fresh blood was divided in 1 to 2 mL aliquots in 14 mL polypropylene tubes with round bottom, pre-warmed for 30 minutes at 37°C under agitation at 180 rotation per minute (rpm), then inoculated with 1/10 (100 μL to 200 μL) of the prepared bacteria to reach a concentration of around 2×10^7^ CFU/mL. Blood was incubated at 37°C under agitation at 180 rpm, and bacteria were enumerated at different time points from 0 to 24 hours post-inoculation. At each time point, blood was gently mixed by slowly pipetting with a P1000 pipette, and 20 μL were added to 180 μL PBS for bacterial enumeration. For serum survival, bacterial growth curves were performed in 10 mL of LB supplemented with 2.5mM CaCl_2_ or in 10 mL of human serum at 180 rpm in 125 mL polycarbonate flasks. Serum inactivation was performed in a water bath at 56°C for 30 minutes.

### Growth curve, *in vitro* stresses and minimal inhibitory concentration assays

For all experiments, parental CO92 and mutant strain glycerol stocks were thawed from −80°C in LB and grown overnight at 28°C under agitation

For the growth curves, bacterial concentrations were adjusted in LB media and seeded at OD_600_ 0.05 in 96-well plates (flat bottom), then incubated at 28°C and 37°C under orbital shaking amplitude 3 in a Tecan Infinite® M Nano plate reader.

For H_2_O_2_ survival assay, bacteria were subcultured in LB at 37°C for 10 hours and harvested at OD around 0.1-0.2. Bacterial concentrations were adjusted in cation-adjusted Mueller-Hinton (MH II) media and seeded in 96-well plates (flat bottom) at 10^6^ CFU/well with pre-warmed MH II containing varying concentration of H_2_O_2_ in 200µL total volume. Plates were incubated for 24h at 37°C with agitation at 1000 rpm in a ThermoMixer® (Eppendorf). After homogenization by pipetting, 20 µL aliquot were serially diluted and plated for bacterial enumeration at different timepoints.

For neutrophil product survival assays, neutrophils were isolated from EDTA-anticoagulated blood drawn by EFS the same day, using the EasySep™ Direct Human Neutrophil Isolation Kit (STEMCELL Technologies) following manufacturer protocol. Purified cells were resuspended in Hank’s Balanced Salt Solution (HBSS) with Ca^2+^ and Mg^2+^ and stimulated 4 hours at 37°C 5% CO_2_ with 25 nM of protein kinase C activator phorbol 12-myristate 13-acetate (PMA, inducing netosis), 5 µM of the ionophore nigericin for degranulation, or left untreated in HBSS. Supernatants were collected after cell centrifugation at 1,000 x g for 10 minutes at 4°C, and kept at −80°C. After overnight growth (see above), bacteria were subcultured in LB or MH II at 37°C for 10 hours and harvested at OD around 0.4. 1 to 2×10^6^ bacteria were harvested by centrifugation, resuspended in 200 µL neutrophil supernatant or HBSS and incubated at 37°C under agitation in a 96-well plate (flat bottom) or 1.5 mL tubes in a ThermoMixer® (Eppendorf). After homogenization by pipetting, 20 µL aliquot were serially diluted and plated for bacterial enumeration at different timepoints.

For the minimal inhibitory concentration (MIC) assays, parental CO92 and mutant strain glycerol stocks were thawed from −80°C in LB and grown overnight at 28°C under agitation. Bacteria were then subcultured in LB at 28°C or 37°C for 10 hours and harvested at OD around 0.4. Bacterial concentrations were adjusted in MH II media and seeded in 96-well plates (round bottom) at 0.5-1×10^6^ CFU/well with pre-warmed MH II containing varying concentration of PmB or LL-37. Plates were incubated for 48h at 28°C or 37°C before picture acquisition of the plate bottom.

### Macrophages intracellular survival

Human blood derived macrophages were isolated and differentiated from buffy coats. Briefly, blood mononuclear cells were isolated by density gradient centrifugation (Ficoll-Paque Premium, Sigma Aldrich, St. Louis, MI, USA). Monocytes were purified from peripheral blood mononuclear cells by positive selection with magnetic CD14 MicroBeads using LS columns (Miltenyi Biotec, Bergisch Gladbach, Germany). Monocytes were cultured for 7 days at 37°C and 5% CO_2_ in Roswell Park Memorial Institute (RPMI) 1640 supplemented with 10% heat-inactivated fetal bovine serum (FBS), L-glutamine and M-CSF (20 ng/mL; R&D systems). Cells were fed every 2 days with complete medium supplemented with M-CSF. RAW 264.7 macrophages from ATCC (TIB-71) were prepared in RPMI with 10% fetal bovine serum with or without IFNγ. Cells were seeded at around 10^4^ cells/well in 96-well plates (flat bottom) and incubated overnight at 37°C 5% CO_2_. Bacteria were grown in 10 mL LB at 28°C under agitation at 180 rpm to reach an OD_600_ of 0.4. Bacteria were serially diluted to reach around 5-10×10^5^ CFU/mL in complete cell media. After removing of the old cell media, 100 µL of the inoculum suspension were added to the cells. The plates were centrifugated at 188 x g (1,000 rpm) for 5 min then incubated 1 hour at 37°C 5% CO_2_. Supernatant was then discarded, and wells were washed three times with cell culture media. Cells were washed once with cell media supplemented with 10 µg/mL gentamycin, then 100 µL of cell media with gentamycin were added to the wells. Plates were incubated for 1h at 37°C 5% CO_2_ until the first timepoint (2 hours) or longer for the next timepoints. After incubation, wells were washed three times with cell media without gentamycin, 100 µL of water were added to the wells and well bottoms were scratched to lyse the cells. For enumeration, inoculum and timepoint were serial diluted and plated on LB hemin at 28°C for 48 hours.

### Neutrophil intra- and extracellular survival

Neutrophils were isolated from fresh EDTA-anticoagulated whole blood one hour after blood draw from a healthy donor using MACSxpress® Whole Blood neutrophil isolation kit (Miltenyi Biotec, Bergisch Gladbach, Germany) following manufacturer instruction. After isolation of neutrophils, red blood cells were depleted by negative magnetic selection with CD235a (Glycophorin) MicroBeads using LS columns (Miltenyi Biotec, Bergisch Gladbach, Germany). Purified neutrophils were resuspended in RPMI 1640 supplemented with 10% heat-inactivated FBS and 10 mM 4-(2-hydroxyethyl)-1-piperazineethanesulfonic acid (HEPES) and dispatched in 96-well plates at 2×10^5^ cells/well. Parental CO92 and mutant strain glycerol stocks were thawed from −80°C in LB and grown overnight at 28°C under agitation, then plated and incubated for 24 hours on LBH plates at 25-28°C. Bacteria were resuspended in 3 mL PBS, vortexed for 1 minute and the OD_600_ was adjusted to 0.25 in 10 mL PBS to reach a concentration around 2×10^8^ CFU/mL, then diluted ¼ in complete cell media. 100 μL of bacterial suspension were added on cells at a multiplicity of infection of 25. Plates were centrifugated for 10 minutes at 2000 rpm and incubated for 1 hour at 37°C with 5% CO_2_. Cells were washed 100 µL complete cell media, then fresh cell medium with or without 20 μg/mL gentamicin was added, then incubated for 1 and 2.5 additional hours. The medium was then removed and the cell lysed by addition of 100 μL of H_2_O for 10 minutes at room temperature. Lysates were serially diluted in PBS to enumerate intracellular bacteria.

### Flea co-infection and proventriculus colonization analysis

Flea infections were performed as previously described^89^. Starved *Xenopsylla cheopis* fleas were allowed to feed for 1 hour on heparinized mouse blood containing 5 × 10^8^ bacteria/mL (cultured overnight at 37°C in BHI) using an artificial feeding system. For co-infection assays, equal amounts of each strain (CO92 pYV- vs CO92 pYV- *slyB*::*dfrB*) were mixed in the blood prior to feeding. Cohorts of 60 female fleas were then collected. Fleas were subsequently fed on mouse blood at 2, 6, 9, and 13-days post-infection.

To assess *Y. pestis* competitive fitness, groups of 20 female fleas were collected at the indicated time points. After decontamination by sequential treatment with hydrogen peroxide and ethanol, individual fleas were placed in 1 mL of PBS and lysed using a FastPrep instrument with Lysing Matrix H (MP Biomedicals). Genomic DNA was then extracted using the Macherey-Nagel Tissue kit according to the manufacturer’s instructions. The total chromosomal copy number of all strains was determined using *proC*-specific primers^90^ (TAAAGCCCCAGTTAATGGCCGATG / ACCAACTTATCGCTAAAATCGACCTG), while the specific abundance of the wild-type strain was quantified using a *slyB* gene-specific primer set (GCTAACAACAACACCTTATCAGG/ ACGCTGAACTGAGTTGGG). Quantitative PCR was performed using Takyon™ ROX SYBR® MasterMix dTTP Blue (Eurogentec) on a Stratagene MxPro 3000 system, following the manufacturer’s recommended cycling conditions. For each sample, *proC* amplification was performed using 1:2 diluted gDNA, whereas *dfr* amplification was performed using undiluted gDNA. DNA copy numbers were used to estimate the total bacterial load (*proC*) and the relative abundance of the wild-type strain (*slyB*) within the population.

To evaluate proventriculus colonization, female fleas were infected with *Y. pestis* expressing green fluorescent protein (GFP) from the pAcGFP plasmid (Addgene). When required, five fleas were randomly selected and dissected under a stereomicroscope to isolate the digestive tract. Immediately after dissection, fluorescence images were acquired using an Eclipse Ci-S fluorescence microscope (Nikon) equipped with a DS-Fi1 camera (Nikon). Images were processed using ImageJ^91^. Briefly, the blue channel was extracted, and an intensity threshold was applied to minimize tissue autofluorescence while preserving the GFP signal.

### Biofilm assay

Biofilm formation was assessed as previously described^21^. Briefly, bacteria (1 × 10⁷ cells in 1 mL) were cultured in LB supplemented with 4 mM MgCl_2_ and 4 mM CaCl_2_ in 24-well plates for 48 hours at 21°C with shaking (180 rpm). The medium was then removed, and adherent biofilms were stained with 0.01% crystal violet for 15 min at room temperature. Wells were rinsed three times with distilled water, and the bound dye was solubilized using an ethanol-acetone solution (80:20, v/v). The absorbance was measured at 542 nm using a Victor X3 plate reader (PerkinElmer).

### c-di-GMP quantification

Intracellular c-di-GMP levels were measured using the CensYBL biosensor as previously described^92^. Briefly, bacteria were transformed with a plasmid encoding split YPet fragments fused to the C-terminal domain of BldD, enabling fluorescence complementation upon c-di-GMP-dependent dimerization. Strains were grown overnight in BHI, diluted to an OD_600_ of 0.5, and incubated under inducing conditions (IPTG) for 2 hours. Cells were then washed in PBS, and fluorescence was measured using a Victor X3 plate reader (PerkinElmer). YFP fluorescence (excitation 490 nm, emission 535 nm, 0.4 s integration time) was normalized to mCherry fluorescence (excitation 530 nm, emission 580 nm, 0.1 s integration time) to account for differences in expression levels.

### PAGE and western blotting

Parental CO92 and mutant strain glycerol stocks were thawed from −80°C in LB and grown overnight at 28°C under agitation. For western blotting of unstressed bacteria, bacteria were subcultured in 100 mL LB in 250 mL polycarbonate flasks at 28°C or 37°C for 7 hours, then harvested by centrifugation at 4°C and bacterial pellet were kept at −20°C. For SlyB induction and oligomerization assays, bacteria were subcultured in 25 mL MH II in 100 mL glass flasks at 28°C or 37°C for 10 hours. PmB stress (10 µg/mL) was added for 1 hour, then bacteria were harvested by centrifugation at 4°C and bacterial pellet were kept at −20°C.

For SDS-PAGE analysis, bacterial pellets were resuspended in Laemmli buffer + β-mercaptoethanol and heated at 95°C for 10 minutes. For semi-native polyacrylamid gel electrophoresis (snPAGE), bacterial pellets were resuspended in Laemmli buffer and heated at 95°C for 10 minutes (denaturing condition) or immediately loaded into the gel (semi-native condition). Samples were loaded and proteins separated on a 4-12% polyacrylamid gel (Invitrogen) at room temperature for SDS-PAGE or on ice for snPAGE, then transferred on a PVDF membrane using the iBlot™ transfer device (Invitrogen). Membranes were blocked in Tris-buffered saline buffer (TBS) with 0.1 % Tween 20 (TBST) and 5% milk for 1 hour, then washed with TBST and incubated with primary antibodies in TBST + 1% milk for 1 hour at room temperature (RT). Membranes were washed with TBST, incubated with secondary antibodies for 1 hour at RT, washed with TBST then revealed using SuperSignal™ West Femto (Thermo Scientific™). Membrane or gel was finally stained with PageBlue (Thermo Scientific™).

## Acknowledgements

We would like to thanks Caroline Manet for her advice on Collaborative Cross collection, Vanessa Kremer for her help with the *in vitro* neutrophil experiments, Mariette Matondo, Thibaut Douché, François Becher and Rita Azevedo for the LC-MS/MS-based proteomics, as well as Eric Krukonis for the kind gift of anti-Ail antibody. We are grateful to all members of the *Yersinia* research unit and the French national reference center for plague and other yersiniosis for insightful discussions. We are grateful to the healthy volunteers of the COSIPOP cohort who agreed to the scientific use of their samples and data and to the CRBIP unit CHIP for providing fit-for-purpose biological resources and associated services. The COSIPOP study group is constituted by members who designed and manage the COSIPOP cohort from the Institut Pasteur with the following persons: AROWAS Laurence, CLEMENT Nathalie, DELHAYE Maurine, EL BSAT Mira, FANAUD Christine, NGUYEN Emilie, NOURY Clémence, PERLAZA Blanca-Liliana, VOGTENSPERGER Marie, WASSILA Smail, ZAYOUD Ayla and lastly JOLLY Nathalie and LAUDE Hélène, as co-senior investigators.

This work was funded by Institut Pasteur, Direction Générale de l’Armement, Agence de l’Innovation de Défense, Fondation pour la Recherche Médicale (FDT20220401-711 5222), the Inception program (Investissement d’Avenir grant ANR-16-CONV-0005), ANRS Emerging Infectious Disease (ANRS0349b, ANRS0350b), Université Paris Cité and the French government managed by ANRS-MIE under the France 2030 program (ANRS-23-PEPR-MIE 0002 – DEBS Plague). The *Yersinia* Research Unit is a member of the LabEX IBEID (ANR-10LBX-62-IBEID).

## Declaration of generative AI and AI-assisted technologies in the manuscript preparation process

During the preparation of this work the authors used the LibreChat plateform internal to Institut Pasteur to reformulate and synthetize text written by the authors. After using this tool/service, the authors reviewed and edited the content as needed and take full responsibility for the content of the published article.

## Supplementary Figures

**Supplementary Figure 1.**
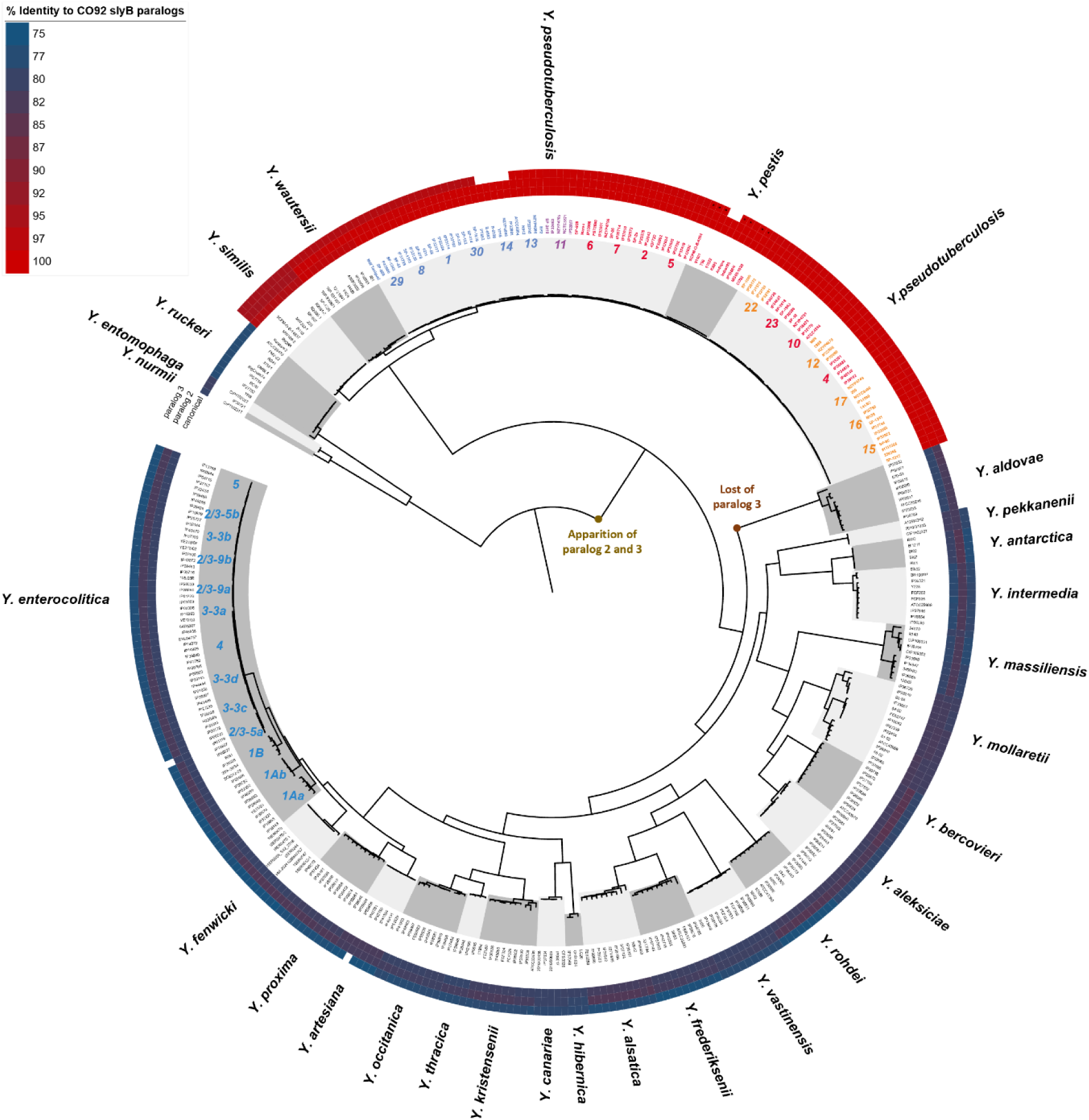
Phylogenetic tree of the *Yersinia* genus showing the presence and homology of *slyB* paralogs compared to *Y. pestis* CO92 alleles. External circles show absence (white) or presence (colored) of the paralog and its nucleotide percentage identity with *Y. pestis* CO92 paralogs. *Y. enterocolitica* and *Y. pseudotuberculosis* genotypes are indicated inside the circles. Strains and *Y. pseudotuberculosis* genotypes are colored according to canonical SlyB allele compared to *Y. pestis* CO92. Red: *Y. pestis* CO92 allele. Orange: A270G synonymous mutation. Purple: G402A synonymous mutation. Blue: G229A non-synonymous mutation (A77T). Frameshifted paralog 3 in *Y. pestis* are indicated with a star inside the corresponding tile. BLAST results can be found in Supplementary Table 1.

**Supplementary Figure 2.**
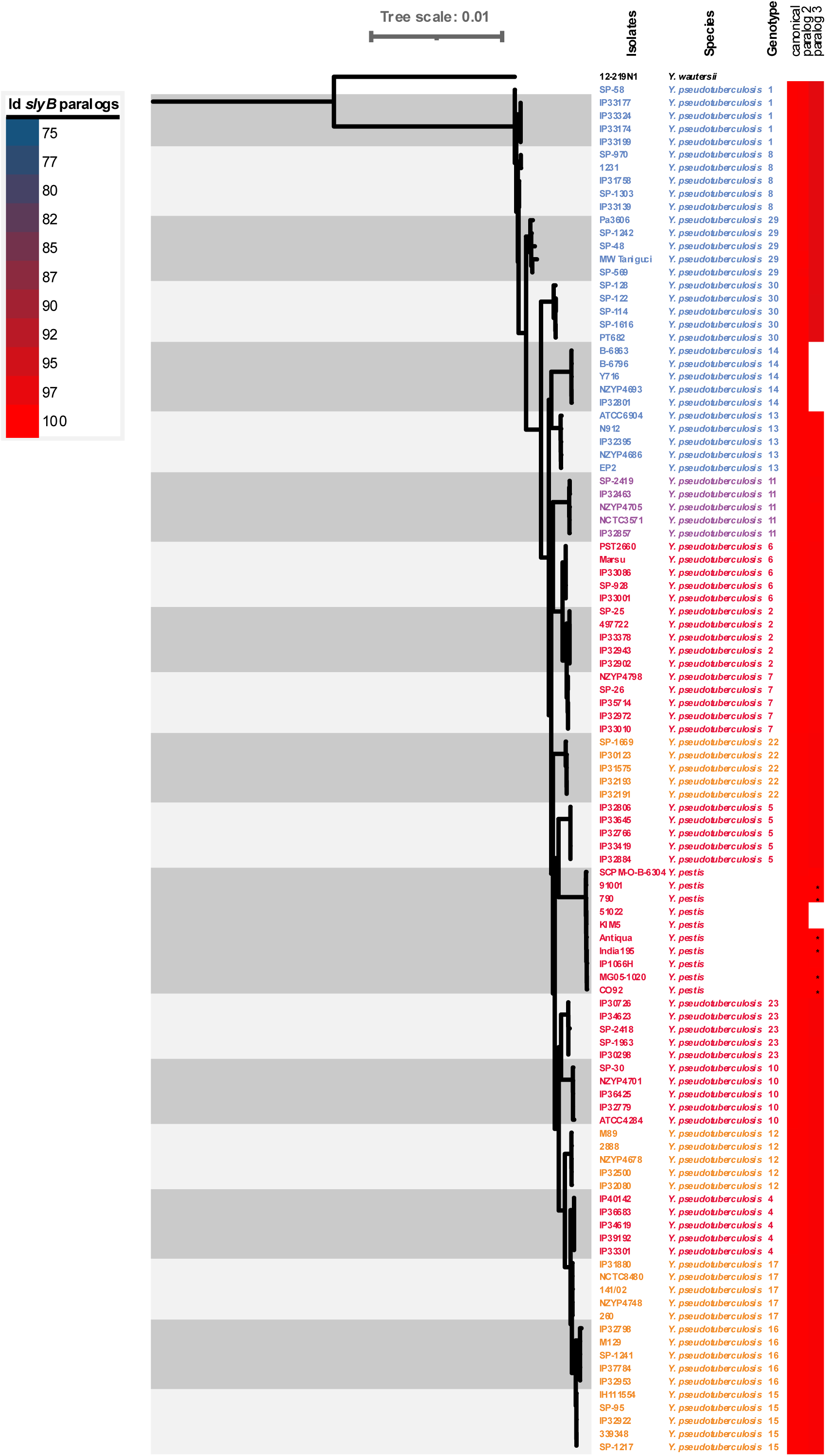
Phylogenetic tree focus from Supplementary Fig. 1 on the *Y. pseudotuberculosis* species. Strains and *Y. pseudotuberculosis* genotypes are colored according to canonical SlyB allele compared to *Y. pestis* CO92. Red: *Y. pestis* CO92 allele. Orange: A270G synonymous mutation. Purple: G402A synonymous mutation. Blue: G229A non-synonymous mutation (A77T). Frameshifted paralog 3 in *Y. pestis* are indicated with a star inside the corresponding tile. BLAST results can be found in Supplementary Table 1.

**Supplementary Figure 3.**
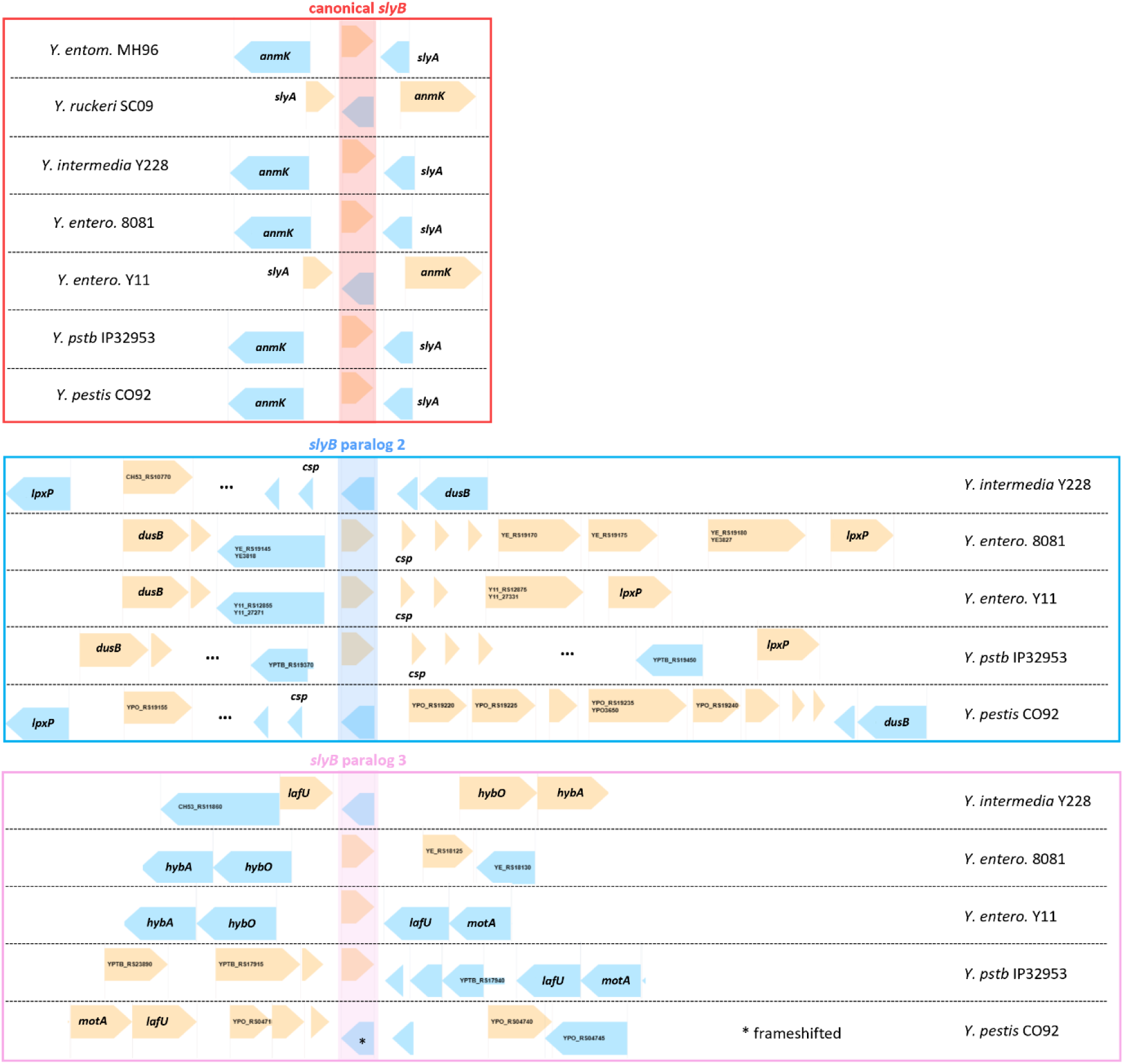
Genetic environment of the 3 *slyB* paralogs in various species from the *Yersinia* genus as shown on the Yersiniomics database. Gene orientation varies depending on the assembly annotation. entom.: *entomophaga*. entero. : *enterocolitica*. pstb: *pseudotuberculosis*.

**Supplementary Figure 4.**
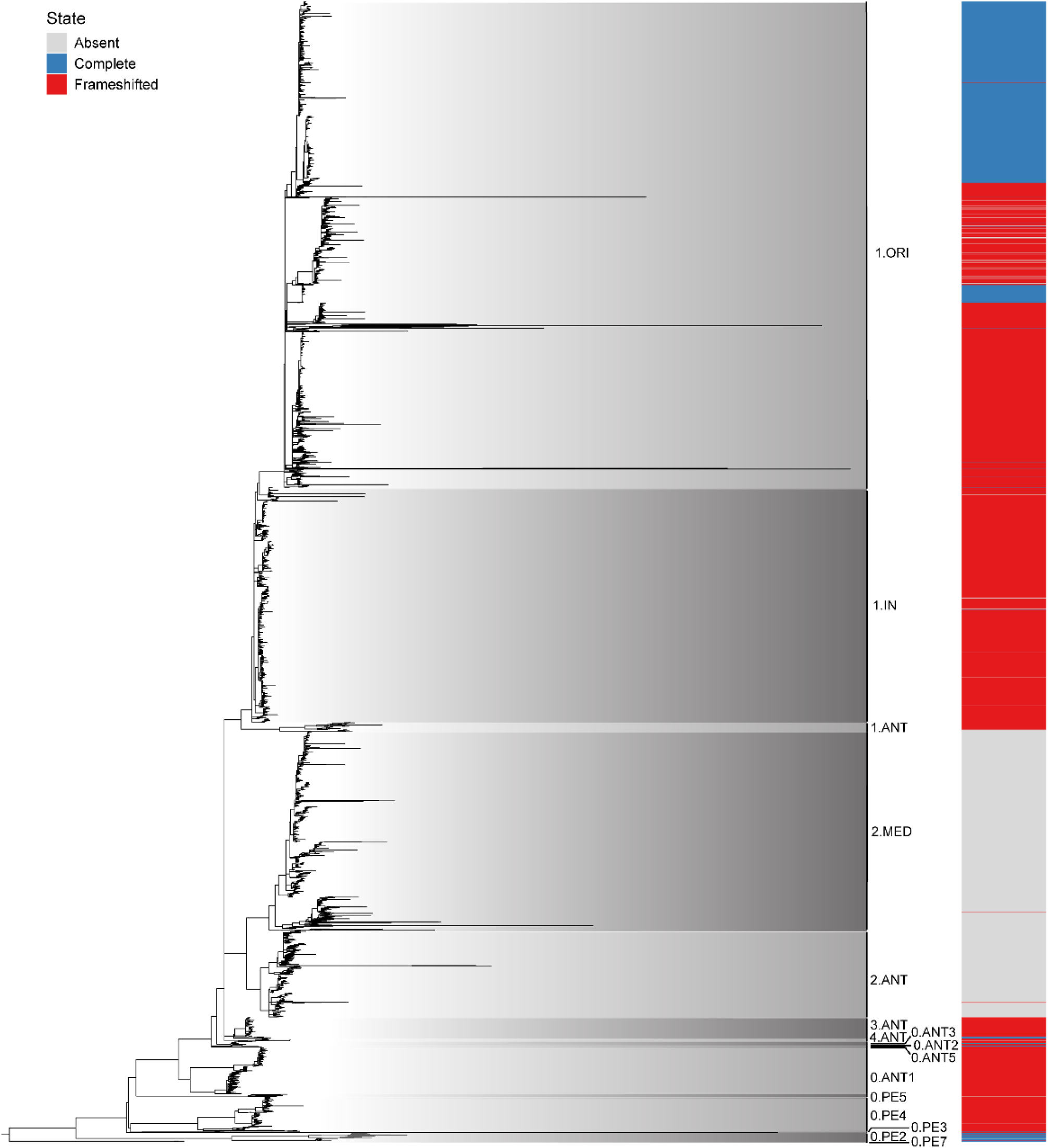
Status of *slyB* paralog 3 across *Y. pestis* evolution. The tree displays the maximum-likelihood phylogeny of 5,744 *Y. pestis* isolates obtained from Mas Fiol *et al.*^46^, with the distribution of 20 major lineages with contemporary isolates labelled next to the tips. The color strip shows the functional status of paralog 3 in each genome, with blue indicating a complete functional sequence, grey indicating absence of sequencing coverage and red indicating a frameshifted version. Detailed results can be found in Supplementary Table 2.

**Supplementary Figure 5.**
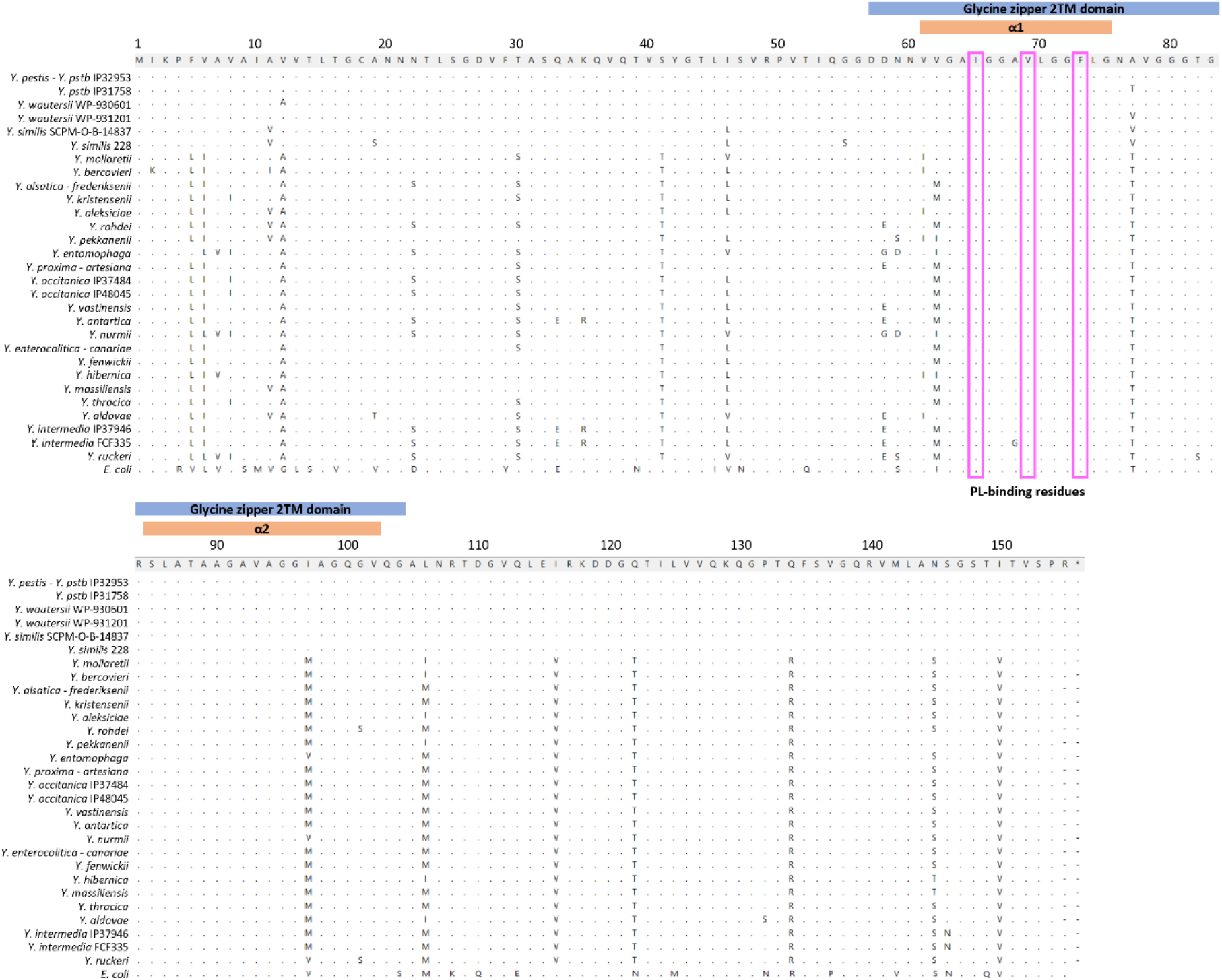
Amino acid (aa) alignments of canonical SlyB from *E. coli* and 28 *Yersinia* species, sorted by lower (top) to higher (bottom) nucleotide distance to *Y. pestis* allele, and showing only aa different from *Y. pestis* allele. Aa numbering is from the 155 aa canonical SlyB*_Ec_* and SlyB*_Yp_*. The glycine zipper 2TM domain and the 2 α-helix containing the glycine zipper are annotated in blue and orange respectively, and the phospholipid (PL)-binding residues are highlighted in pink in the alignments.

**Supplementary Figure 6.**
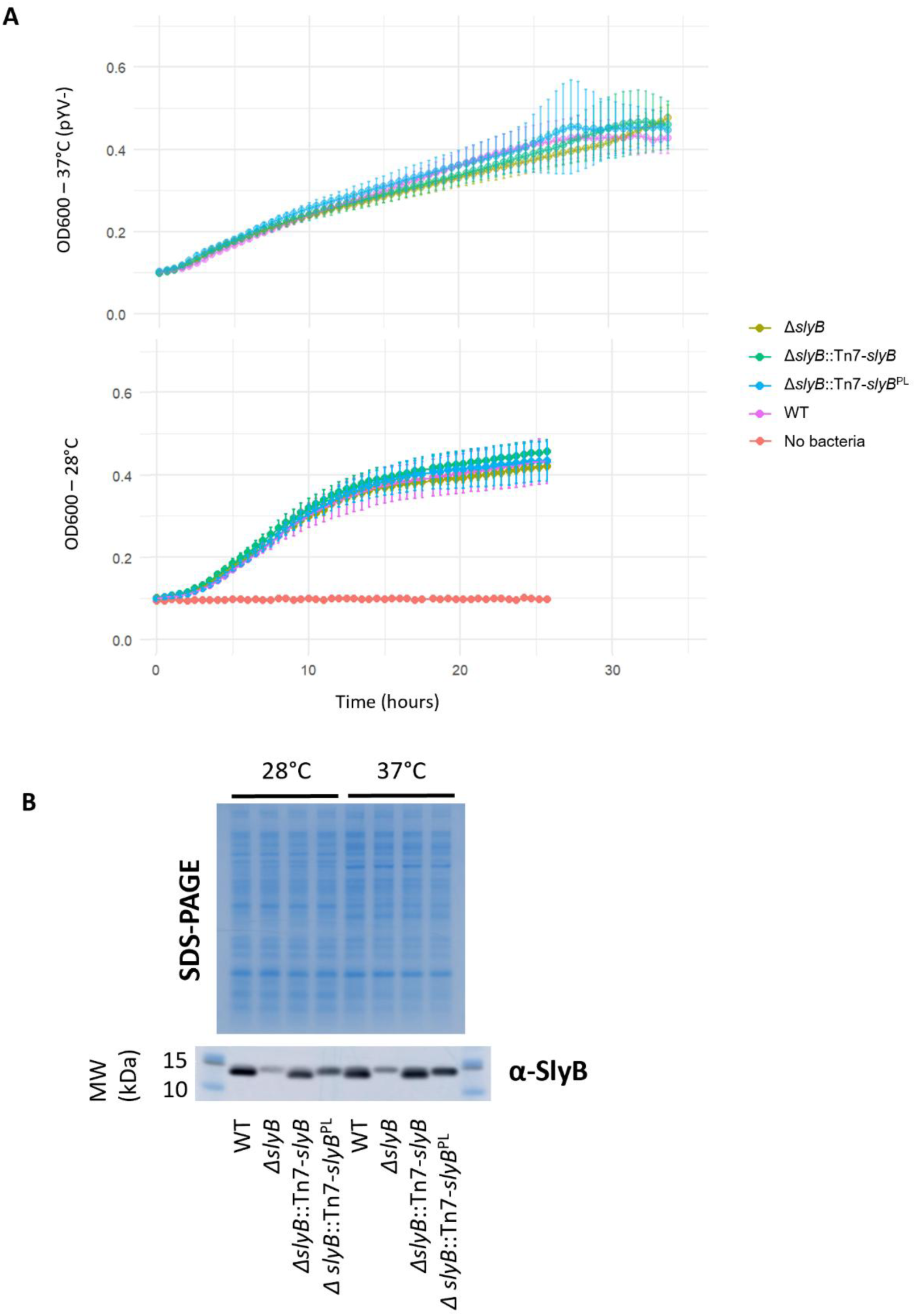
(A) Growth in LB of different *slyB* constructs at 28°C (pYV+) and 37°C (pYV-). (B) SDS-PAGE and associated anti-SlyB*_Ec_* western blot of *Y. pestis* CO92 construct whole cell lysates. Weak expressions of a cross-reacting protein in the Δ*slyB* strain could correspond to the second *slyB* paralog YPO3646. MW: molecular weight.

**Supplementary Figure 7.**
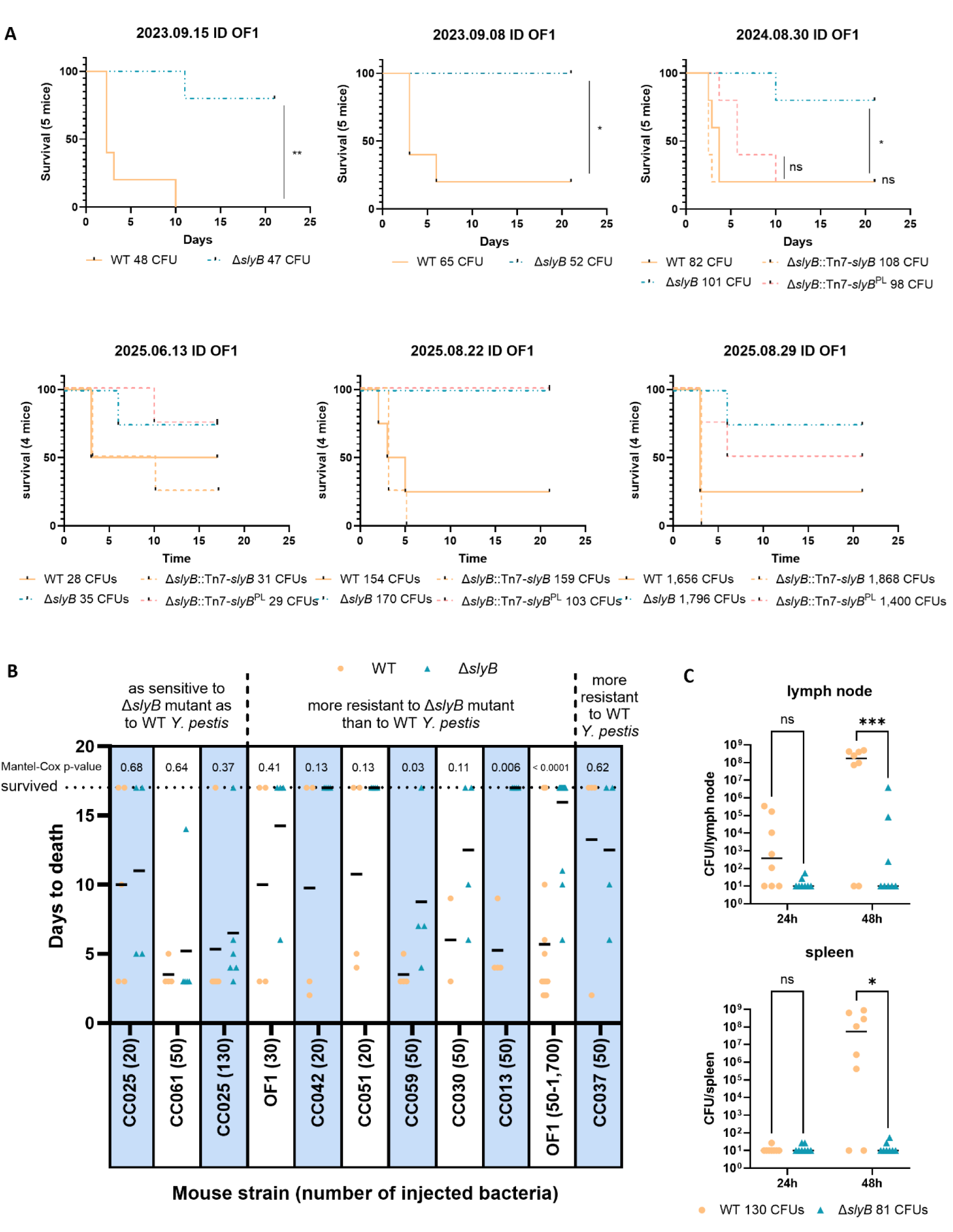
(A) Survival of OF1 mice after intradermal injection in the ear pinna. Data show each experiment performed with different doses on different days. Statistical significancy was assessed by Mantel-Cox logrank test. (B) Survival of 8 Collaborative Cross (CC) inbred lineage and outbread OF1 mice after intradermal injection of WT and Δ*slyB* strains. Estimated injected bacteria is indicated in bracket for each experiment. Statistical significancy was assessed by Mantel-Cox logrank test. (C) Bacterial load in OF1 draining lymph node and spleen after intradermal injection in the ear pinna of approximatively 130 and 81 CFUs of the WT and Δ*slyB* constructs. Šidák’s multiple comparisons test was used for statistical testing. ns: not significative. *: p-value < 0.05. **: p-value < 0.01. ***: p-value < 0.001.

**Supplementary Figure 8.**
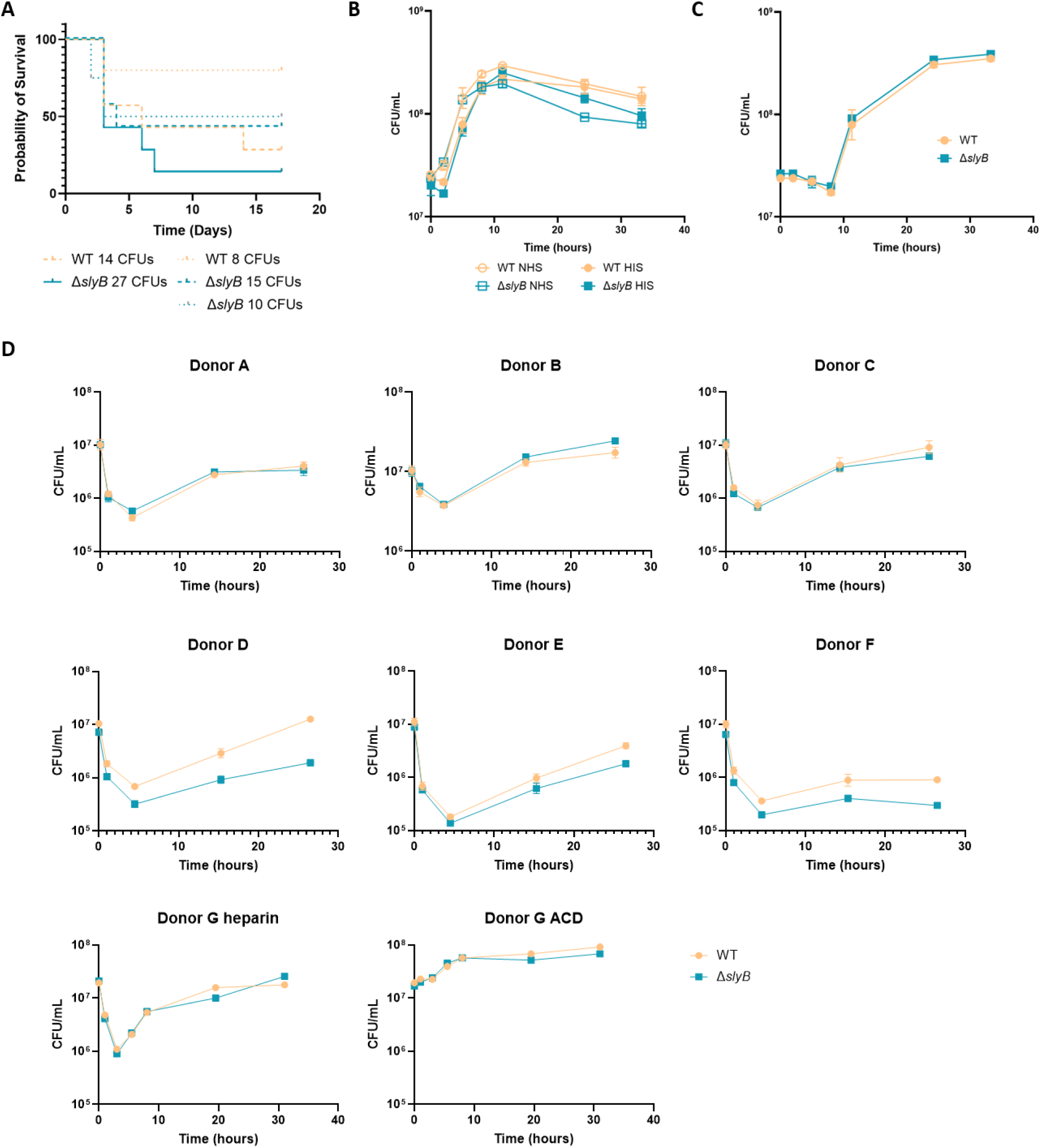
(A) Survival of OF1 mice after intravenous injection in the tail vein. (B) Bacterial growth in normal human serum (NHS) and heat-inactivated serum (HIS) at 37°C. (C) Bacterial growth in LB supplemented with 2.5 mM Ca^2+^ at 37°C. (D) Bacterial survival in human whole blood anticoagulated with heparin (donors A to F) or comparing heparin and citrate-dextrose solution (ACD) anticoagulant (donor G).

**Supplementary Figure 9.**
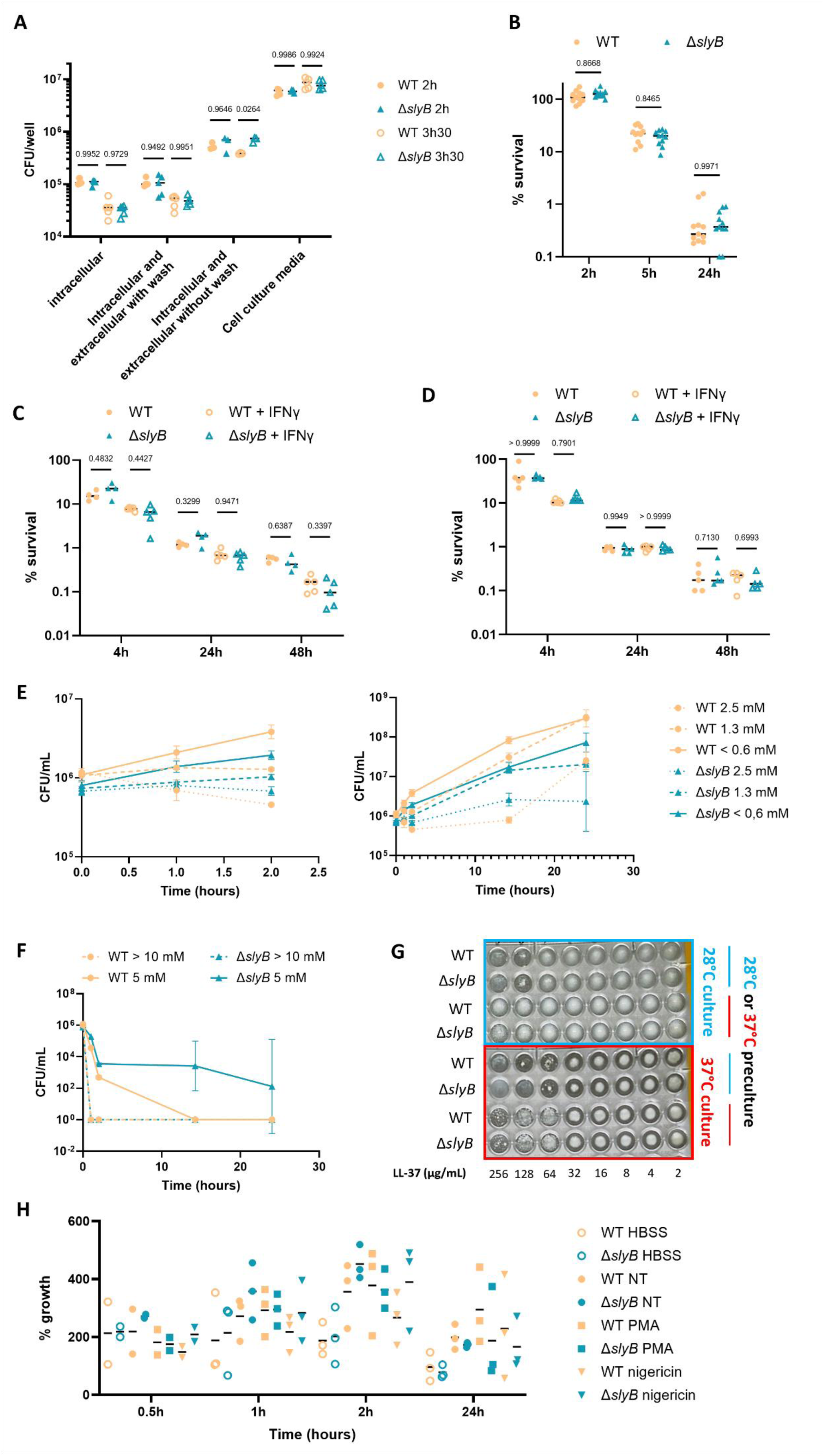
(A) Bacterial survival of pYV+ constructs upon exposure to human neutrophils. Intracellular survival was assessed by addition of gentamicin. Extracellular and intracellular survival was assessed without gentamicin treatment and with or without well wash with PBS. (B) Intracellular survival of pYV+ constructs in human blood monocyte-derived macrophages. (C-D) Intracellular survival of pYV+ constructs in RAW 264.7 macrophages activated or not with IFNγ with MOI around 6 (C) or 4 (D). Survival is expressed as percentage of CFUs compared to the inoculum. Statistics were computed on log-transformed value by two-way ANOVA with Šidák’s multiple comparisons test. (E-F) Survival of pYV- constructs upon exposure to H_2_O_2_ in MH II media at 37°C up to 24h, with concentration below (E) and above (F) 5mM. (G) Survival of pYV+ constructs upon exposure to the cAMP LL-37 in MH II media at 28°C or 37°C for 48h. (H) Survival of pYV- constructs upon incubation at 37°C in Hank’s Balanced Salt Solution (HBSS) or supernatant of neutrophils stimulated with the protein kinase C activator phorbol 12-myristate 13-acetate (PMA), the ionophore nigericin or untreated (NT). Growth is expressed as percentage of CFUs compared to T0.

**Supplementary Figure 10.**
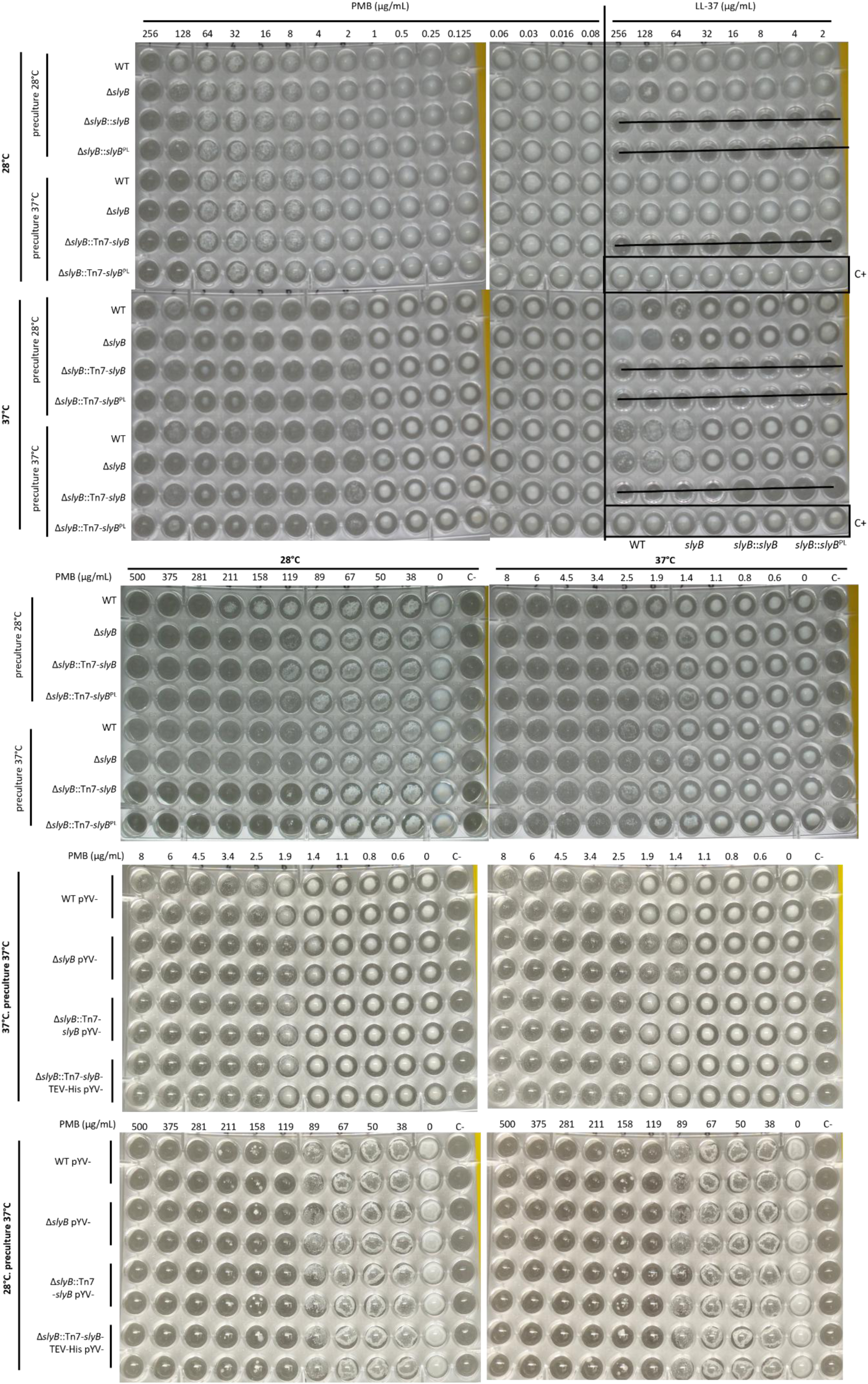
Polymyxin B (PmB) and LL-37 minimum inhibitory concentration (MIC) for diverse *slyB* constructs. After preculture at mid-exponential phase in LB at 28°C or 37°C, PmB (or LL-37) exposition was done for 48h at 28°C and 37°C in cation-adjusted Mueller-Hinton (MH II) media.

**Supplementary Figure 11.**
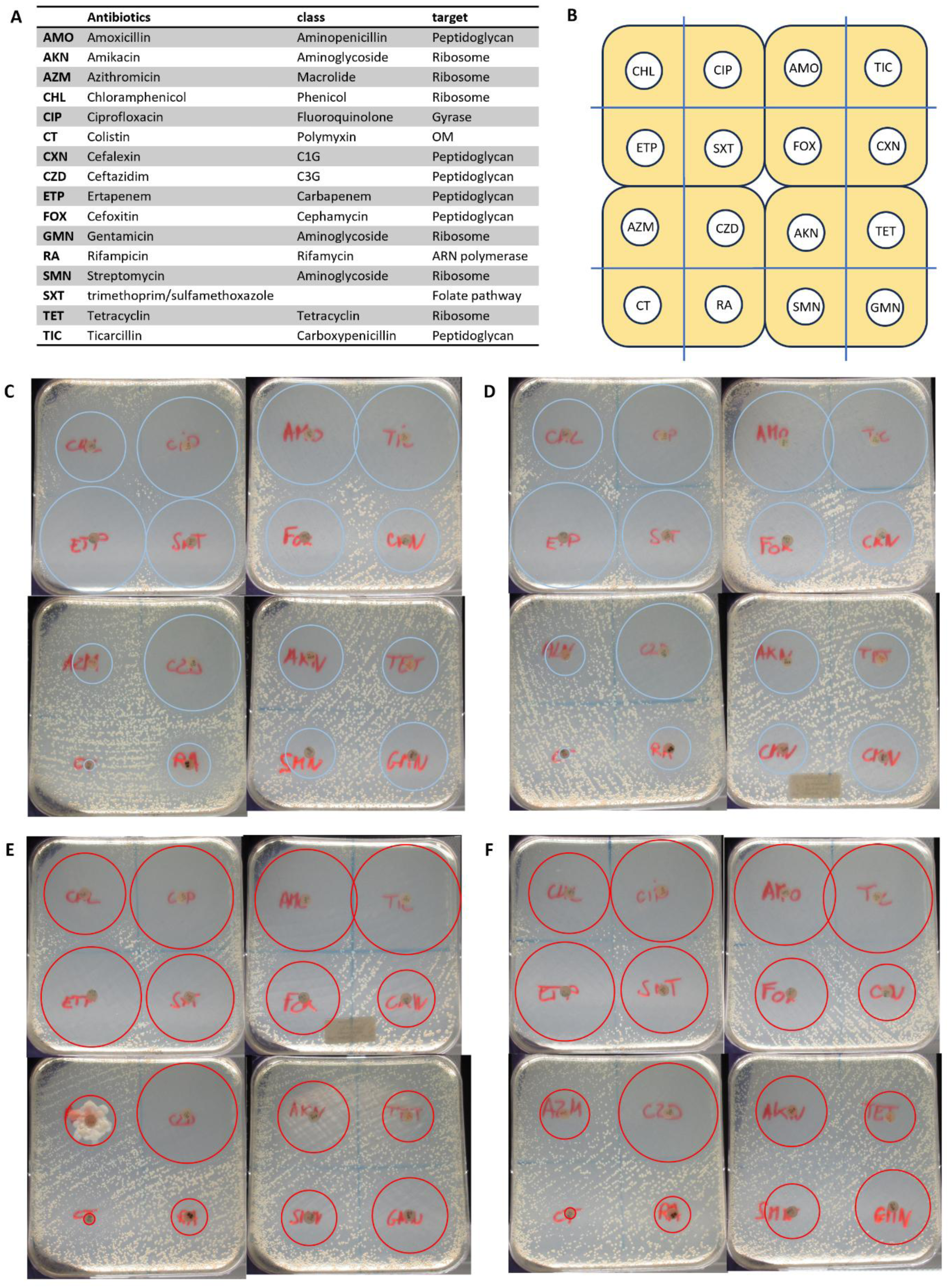
Disk diffusion assays on 12 antibiotics on MH II agar plates. (A) List of antibiotics used, with associated class and molecular target. (B) Disk dispositions on plates for the disk diffussion assay. (C-F) Disk diffusion assay for (C and E) WT and (D and F) Δ*slyB* strain at (C and D) 28°C and (E and F) 37°C Inhibition area is drawn in blue or red. OM: outer membrane.

